# Native-State Imaging Reveals Spatially Separated Organized Cells and Strain-Specific Matrix Architecture in *Pseudomonas aeruginosa* Biofilms

**DOI:** 10.1101/2025.08.28.672782

**Authors:** Goodness O. Osondu-Chuka, Guruprakash Subbiahdoss, Stephan Schandl, Aleksandr Ovsianikov, Olivier Guillaume, Erik Reimhult

## Abstract

*Pseudomonas aeruginosa* (PA) biofilms resist antibiotics and immune clearance through their multicellular community organization. Yet, the native spatial arrangement of cells and the extracellular matrix (ECM) remain poorly understood, as research has largely focused on genetics and regulatory networks rather than physical structure. This has limited our understanding of how physical structural organization connects to biofilm function. We combine high-pressure-freezing cryo-SEM, confocal microscopy, and quantitative spatial analysis to examine 4-day-old mucoid and PAO1 biofilms in a near-native state. In contrast to the dense cellular aggregates often inferred from dehydrated samples, cryo-SEM revealed bacterial cells as individually embedded within a continuous extracellular matrix. Spatial statistics revealed a preferred intercellular spacing of approximately 1 µm, broad spacing distributions, and only weak short–range clustering. Depth-resolved confocal analysis confirmed this sparse organization across larger biofilm volumes and revealed vertical stratification, with the highest bacterial volume fraction and a lower cell volume fraction but stronger clustering toward the biofilm surface. Despite these common structural features, local orientational order exhibited significantly different depth-dependent profiles between the two phenotypes, tending toward higher order at the biofilm base in mucoid biofilms and at the biofilm surface in PAO1. Cryo-SEM further showed that mucoid biofilms contained aligned fibrillar structures within the biofilm interior, whereas PAO1 biofilms exhibited a denser, mesh-like matrix. These findings challenge prevailing views of biofilms as densely packed bacterial aggregates and establish a quantitative framework for understanding antimicrobial tolerance, cell–cell interactions, nutrient access, and biofilm mechanics in both clinically and environmentally relevant contexts.

## 1 Introduction

*Pseudomonas aeruginosa* (PA) is a highly adaptable, opportunistic pathogen known for forming robust biofilms that contribute to persistent infections and antibiotic resistance.[1–3] Central to this resilience is the biofilm’s extracellular matrix (ECM), a complex network of polysaccharides (alginate, Pel, and Psl), proteins, and extracellular DNA (eDNA) that acts as a robust protective barrier, enabling bacteria to withstand high antibiotic concentrations.[4,5] Alginate, in particular, is a hallmark of mucoid clinical isolates, often found in cystic fibrosis (CF) lungs, where it enhances biofilm resilience.[4,6–8] This resilience is primarily attributed to its ability to provide structural stability and act as a physical barrier against both antibiotics and host immune defenses.[4,6,8] Overproduction of alginate leads to biofilms that are notably persistent and difficult to eradicate in clinical settings, making it a critical target for research.[6,7] Understanding the structural complexity of these biofilms is therefore essential for developing effective therapeutic strategies, as PA biofilm formation plays a central role in chronic infections, such as those associated with CF, burn wounds, and indwelling medical devices.[4,9]

Most of our understanding of biofilm formation and structure comes from detailed studies of a few species and strains, with *P. aeruginosa* PAO1 perhaps the best-studied.[3,10,11] PAO1 (non-mucoid, reference strain) and mucoid PA, despite belonging to the same species, exhibit distinctly different extracellular matrix profiles and form different biofilms in terms of both biochemical composition and large-scale structure.[12,13] These differences may significantly influence bacterial clustering, intercellular spacing, and spatial distribution. A significant gap remains in our understanding of nano- and microscale biofilm structure, including how cells and ultrastructure are organized; we must bridge these scales to understand biofilm function at length scales relevant to bacterial communication and community organization.

Electron microscopy has long been employed in biological research to visualize the ultrastructure of cells, bacteria, and tissues. Among its variants, scanning electron microscopy (SEM) has emerged as a valuable tool for visualizing PA biofilms, providing high-resolution images that reveal intricate surface morphology and spatial organization.[14–16] However, biofilm hydration and complex extracellular matrix composition, particularly with polysaccharide-rich regions,[17–19] pose a significant challenge in electron microscopy, where traditional methods (air-drying, chemical-drying, and critical point drying (CPD)) require sample dehydration for imaging in vacuum, leading to structural collapse and loss of integrity.[20–22] Membranes and polysaccharides undergo significant reorganization, shrinkage, and collapse upon water removal, distorting their native architecture and functional properties.[18,23,24] Cryo-scanning electron microscopy (cryo-SEM) offers a solution for preserving biofilm structures in their near-native state.[20,21] Combined with high-pressure freezing (rapid freezing of samples within milliseconds to prevent ice crystal formation) and freeze-etching, cryo-SEM enables high-resolution visualization of delicate extracellular polymeric substances (EPS), cellular arrangements, and internal biofilm ultrastructure that are lost using conventional preparation methods.[25–27] High-resolution, high-fidelity imaging with cryo-SEM can also quantify these structures within biofilms.[21,28]

Another indispensable tool for investigating biofilm structure and organization in 3D is confocal laser scanning microscopy (CLSM).[29,30] As an ideal complement to cryo-SEM, CLSM enables the visualization of cellular distribution across the biofilm volume, making it particularly well-suited for quantitative structural analysis.[31–33] Unfortunately, its resolution is diffraction-limited, putting ultrastructural analysis out of reach and making analysis of densely packed bacteria in biofilms challenging. However, by applying statistical image analysis tools, bacterial cell organization in biofilms can be quantified and verified with cryo-SEM snapshots. By segmenting 3D CLSM image stacks into defined volumetric elements, it becomes possible to extract layer-resolved metrics describing how cells are distributed and organized along the biofilm thickness.[34,35] Such analyses provide insight into structural anisotropy and vertical heterogeneity, which are critical features of the biofilms but are not readily accessible through using only 2D imaging techniques.[34,35] Beyond spatially quantifying cell volume fraction and clustering, imaging bacterial orientation during biofilm formation can reveal whether their architectures exhibit coordinated local orientational ordering, which may indicate emerging multicellular organization even when cells are not densely packed.

The aim of our work was to elucidate the organization and ultrastructure of PA biofilms in their native state by combining the strengths of cryo-SEM and CLSM for quantitative image analysis. By minimizing preparation-induced artifacts and revealing the native structural organization of cells and ECM, we seek to provide missing structural information beyond genetic and biochemical data, e.g., whether *P. aeruginosa* always organizes into microcolonies and forms dense cellular layers within a homogeneous matrix near the substrate. Such knowledge could prove instrumental in elucidating how biofilms form and grow, establishing the causes of biofilm tolerance to antibiotics, and developing more effective anti-biofilm strategies.

## 2 Results and Discussion

### 2.1 *P. aeruginosa* biofilms exhibit sparse, spatially organized cells rather than dense microcolonies

SEM is a vacuum-based imaging technique, and structural observations of naturally hydrated systems can vary with sample preparation methods, which is highly relevant to its use in biofilm research.[18,36,37] We performed an initial study establishing that high-pressure freezing (HPF), with cryo-SEM imaging, was the only approach, compared with air-drying and critical-point drying (CPD), that could preserve the structure of cell distribution and the extracellular matrix in PA biofilms (cf. **Figures A.4 and A.15**). This result agrees with previous studies demonstrating that air-drying and CPD can cause dehydration-associated EPS collapse and biofilm shrinkage, whereas HPF–cryo-SEM better preserves bacterial ultrastructure and ECM organization.[26,38,39].

To quantify the distinct spatial organization observed in the cryo-prepared PA biofilms, we calculated the centroid nearest-neighbor distances (NND) and the radial distribution function (RDF; *g*(*r*)) for the distribution of cells obtained from multiple cryo-SEM samples (**Figures 1 and A.5**, **respectively**) for different strata within the biofilm: bottom (∼1 – 5 µm closest to the substrate), bulk (∼5 – 45 µm from the substrate), and top ( ∼1 – 5 µm closest to the bulk fluid). RDFs and related pair-correlation methods are increasingly being used to resolve preferred cell spacing and short-range order across spatial scales.[41,42] Although quantitative analysis of cell-scale spacing is increasingly being applied to bacterial biofilms, comparable characterization of single-cell spatial distributions in established *P. aeruginosa* biofilms remains limited.

**Figure 1:**
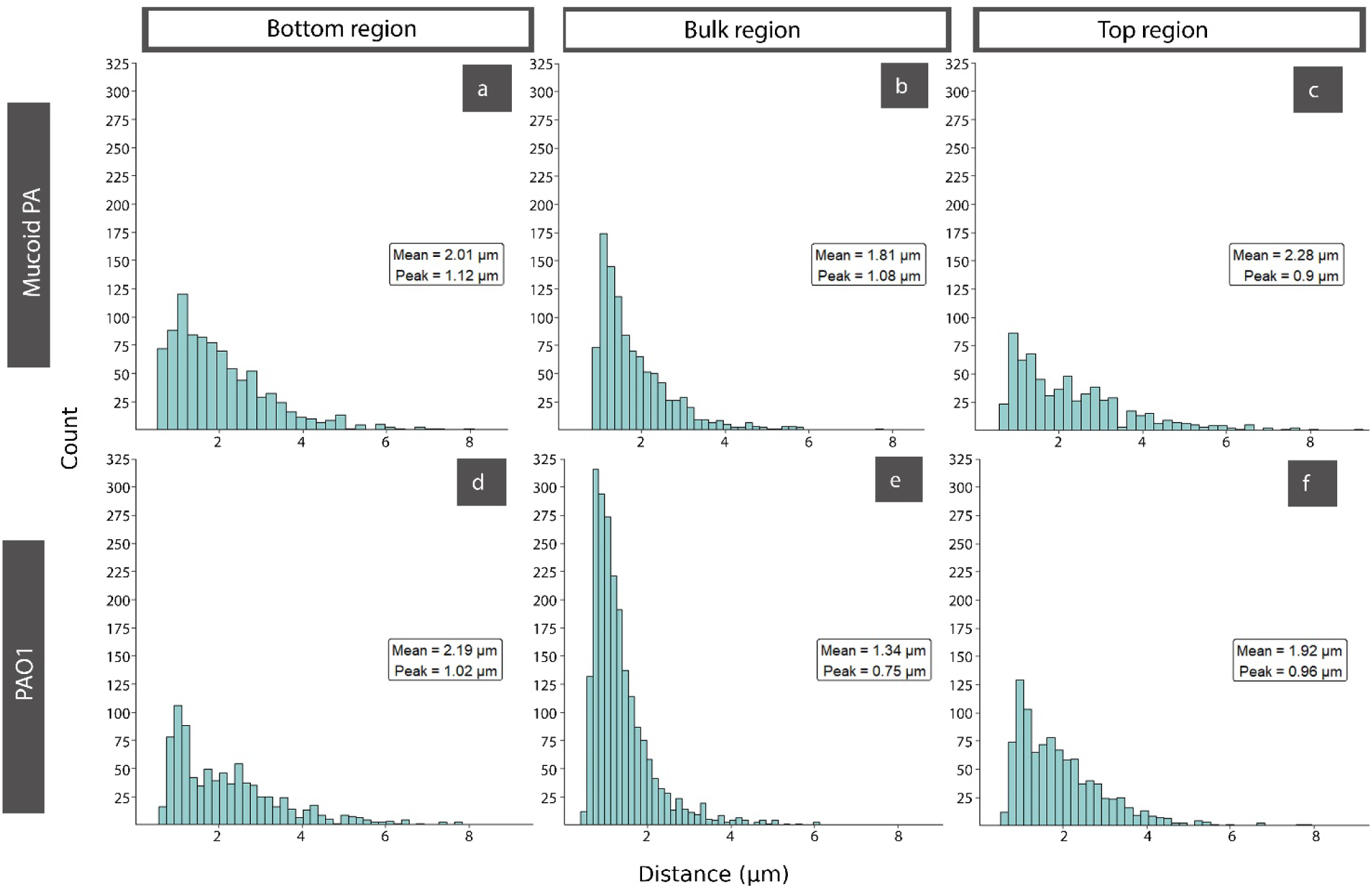
Region-specific distributions of centroid nearest-neighbor distances within PA biofilm. Histograms show nearest-neighbor distances for bacterial cell centroids in (a) bottom, (b) bulk, and (c) top mucoid PA biofilm regions, and (d) bottom, (e) bulk, and (f) top PAO1 biofilm regions.

The NND histograms for both PA biofilms (**Figure 1)** show that although the peak distance is ∼1 µm across strata, the mean distance between neighboring bacteria is ∼2 µm, with a significant tail at distances >4 µm and a broader distribution for bacteria in the top and bottom sections of the biofilms. This is also reflected in larger mean NNDs for the top and bottom. Similarly, the *RDF* analysis showed that direct cell-cell contact is rare, and for the mucoid PA, cells were most likely found at distances of ∼1.1–1.3 µm and multiples thereof, up to a maximum of 4 µm apart. This spatial ordering is much more pronounced for mucoid PA than for PAO1. At long distances, no significant spatial ordering was found. Researchers have used NND and RDF analyses in other bacterial systems to quantify cell-scale spatial organization. For example, mean centroid-to-centroid NNDs of ∼1.1–1.2 µm have been reported in densely populated *Shewanella oneidensis* biofilms, while *Streptococcus mutans* biofilms showed characteristic NND separations of ∼1–2 µm. [35,40,41]

The intercellular spacings observed in this present study suggest that both PA biofilms adopt sparse spatial organizations rather than a purely random distribution of cells or tight clustering. This challenges the classical microcolony-centered model of biofilm development, in which surface-attached founder cells undergo clonal proliferation and local accumulation to form densely packed multicellular structures.[43,44] Instead, it points toward a balance between growth, matrix secretion, and possibly mechanical relaxation processes that collectively regulate cell positioning.[45–47]

These findings have important implications for interpreting biofilm physiology and address a key limitation of dehydration-based imaging approaches, which tend to artificially increase apparent cell clustering and obscure ECM-defined spacing. Dehydration promotes capillary forces and matrix collapse,[38] effectively driving cells into closer contact and inducing clustering that does not reflect their native spatial organization.[38,48] As a result, dried samples may resemble aggregated colloidal systems, potentially leading to an overinterpretation of direct cell–cell interactions. The quantitative demonstration of consistent intercellular distances across biofilm depths provides a more realistic framework for modeling biofilm mechanics and transport processes, particularly at the tens-of-micron length scales that govern diffusive exchange or micron length scales of direct cell–cell force interactions, thereby bridging an important gap between structural observations and functional interpretations in both clinical and environmental biofilms. The observed consistency in preferred distances between strata and across biological replicates supports the conclusion that this architecture is not random packing or a consequence of purely local environmental effects.

### 2.2 Cell clustering in *P. aeruginosa* biofilms is strongest near the biofilm surface and strain-specific

To extend our observations from high-resolution cryo-SEM to assess whether the identified spatial organization persists in larger biovolumes, we employed CLSM-based 3D imaging combined with quantitative spatial analysis. CLSM provides lower spatial resolution but enables analysis of larger volumes and higher numbers of bacteria, thereby improving statistical power. Staining of the bacteria and constraints on cell shape from the cryo-SEM analysis were used to segment the fluorescence and fit prolate *P. aeruginosa* as cylinders, variable in size and orientation (**Figure A.6)**. The centroid coordinates of the cylinder-fitted bacteria were extracted under consistent imaging conditions across >6 biological replicates and used to calculate spatial correlations using Ripley’s *H*-function (*H*(*r*)). Calculation of Ripley’s *H*-function allows us to quantitatively determine and compare between biofilms whether cells are clustered, depleted, or randomly distributed. Importantly, it also shows over which length scales they are organized, which for sparse biofilms, such as mucoid *P. aeruginosa*, can be impossible to infer by eye. Positive *H*(*r*) values indicate clustering relative to a random distribution, whereas values near zero indicate random organization, and negative values indicate fewer bacteria than would be expected by chance. After an initial gap, biofilms of both strains exhibited weakly positive *H*(*r*) values at distances up to ∼8 and ∼6 µm for mucoid PA and PAO1, respectively, across all samples and layers (**Figure 2**), demonstrating that cell distributions in both strains were intrinsically non-random and dominated by local correlations. In both strains, the strongest clustering (pronounced *H*(*r*) occurred at short length scales of approximately 1–2 µm, with the most pronounced clustering observed in the upper biofilm regions. However, differences emerge when clustering behavior is examined across larger spatial scales and along the biofilm depth.

**Figure 2:**
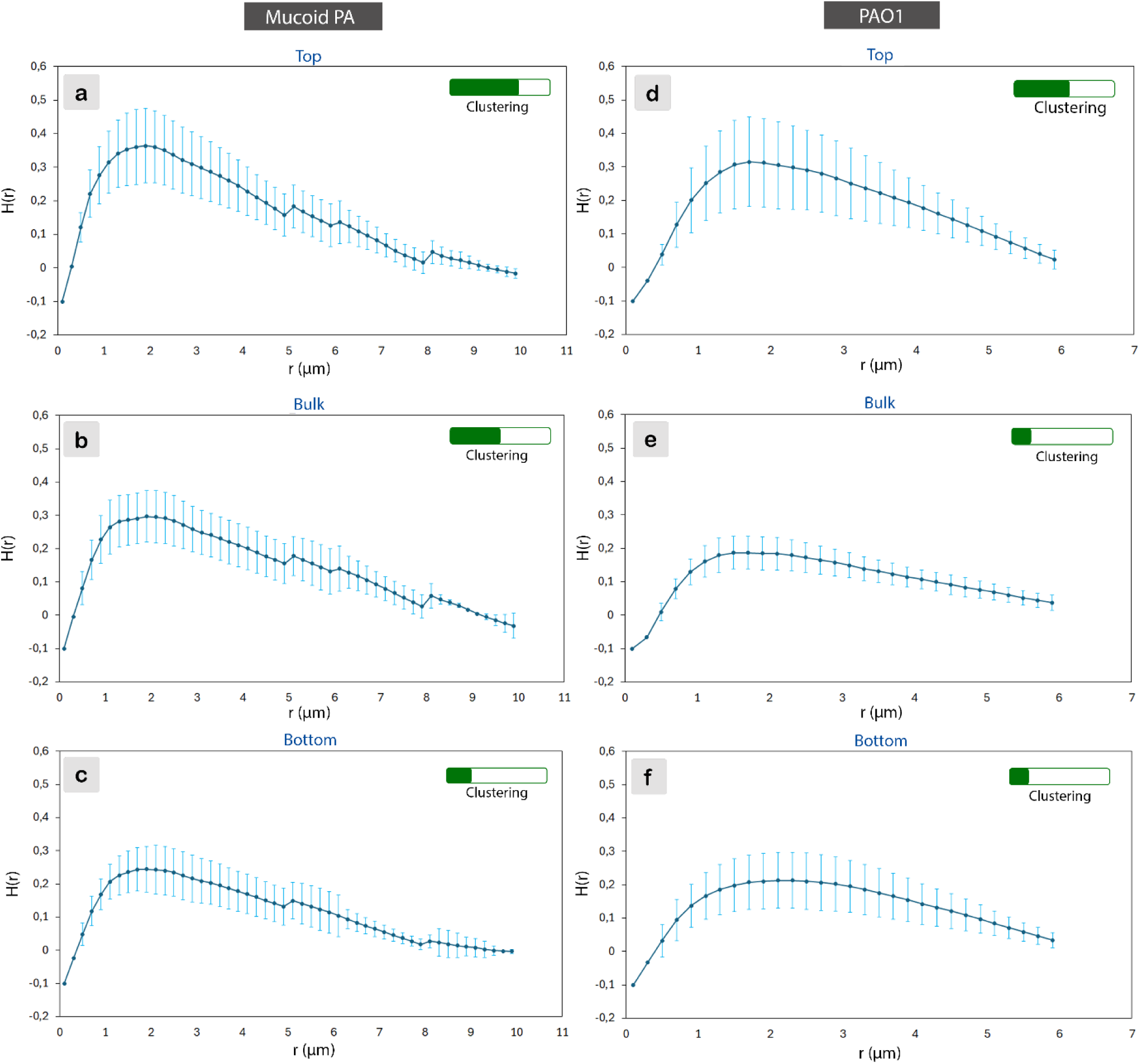
Spatial organization of mucoid PA and PAO1 biofilms revealed by averaged Ripley’s *H*(*r*) analysis. Ripley’s *H*(*r*) functions were calculated for cells located in volumes close to the a,d) biofilm–substrate interface (Bottom), b, e) in the biofilm interior (Bulk), c, f) and close to the biofilm–liquid interface (Top) for mucoid PA (left) and PAO1 (right) biofilms. Curves represent Ripley’s *H*(*r*)values averaged across more than 5 independent biological replicates per strain, and the error bars signify the 95% confidence interval.

Mucoid PA biofilms exhibited vertically stratified spatial organization. Within the top volume, the mucoid strain exhibited the highest degree of clustering, with *H*(*r*) peaking at ∼ 0.38–0.40 at *r* ≈ 1.5 − 2.0 µ*m*, as expected. Positive *H*(*r*) values persisted until *r* ≈ 8 µ*m*, indicating the presence of larger clusters than in the interior of the biofilm. Within the biofilm interior (bulk region), results showed less clustering, with peak *H*(*r*) values of ∼0.30–0.32 at *r* ≈ 1.5 − 2.0 µ*m*. In the bottom volume, mucoid PA biofilms displayed the least clustering. Peak *H*(*r*) values of approximately 0.25–0.26 were observed at *r* ≈ 1.8 − 2.2 µ*m*.

*H*(*r*) values up to 8 µm throughout the mucoid biofilm indicate the presence of clusters, reflecting local correlations in bacterial distribution. However, the cryo-SEM images and *H*(*r*) values of 0.2–0.4 reveal that these clusters are not densely packed cells or microcolonies. CLSM visually exaggerates bacterial packing through background fluorescence and diffraction-limited resolution. A possible interpretation of the moderate overall clustering and the significantly higher clustering at the top of the biofilm is that local clustering in young mucoid PA biofilms primarily reflects the migration of daughter cells from mother cells. Clustering would be more pronounced in a more actively growing layer and less pronounced where cell division has slowed. Presumably, the biofilm grows primarily through the top layer, while cells divide less close to the bottom, leading to the observed clustering behavior, but this hypothesis requires confirmation through live imaging. From a functional perspective, clustering is expected to exacerbate diffusion limitations, promote metabolic stratification, and increase tolerance to antimicrobial treatments, and our results suggest these effects are reduced in rapidly growing young biofilms. [49–51] Similar depth-dependent clustering has been reported in early *Staphylococcus aureus* biofilms; Ripley function analysis revealed progressively stronger clustering with increasing distance from the attachment surface, consistent with the development of vertically organized, column-like structures.[52] However, given the substantial differences in cell morphology, motility, matrix composition, and growth conditions between *S. aureus* and *P. aeruginosa*, the mechanisms underlying these similar spatial patterns likely differ.

PAO1 biofilms exhibited weaker spatial organization and reduced vertical stratification than mucoid PA. Positive *H*(*r*) values declined more rapidly than in the mucoid strain, approaching zero by *r* ≈ 6 µ*m*. The bulk and bottom regions exhibited weaker clustering than the top layer for PAO1 (**Figure 2**), reflecting a more homogeneous internal organization. Hence, while PAO1 shows signs of sparse clusters, these are even less pronounced and smaller than those in the mucoid strain.

Taken together, the cryo-SEM and CLSM analyses consistently demonstrate that cell organization within 4-day-old mucoid and PAO1 biofilms is not random; it is characterized by short-range spatial order, but not by dense cluster or microcolony formation, and by relatively large spacing and a broad distribution of nearest-neighbor spacings. Hence, our findings challenge the conventional view of PA biofilms as tightly packed cell clusters surrounded by ECM, which was derived from observations of collapsed biofilms.[53–56] Evidence from other microbial systems supports the notion that substantial intercellular separation is a genuine feature of hydrated biofilms. Hrubanová et al., for example, used high-pressure freezing and cryo-SEM to preserve fully hydrated *Staphylococcus epidermidis* and *Candida albicans* biofilms and visualized microbial cells separated within the surrounding material rather than uniformly compacted into dense cellular masses.[39] An analogous finding of cell-cell spacing was reported by Kamila *et al.*, who described large distances between microbial cells in cryo-imaged *Staphylococcus epidermidis* and yeast biofilms, but did not analyze it further.[21] Although intercellular spacing was not quantified in that study, these observations are qualitatively consistent with the architecture identified here.

A locally ordered, spaced, and homogeneous distribution of distant bacterial cells in biofilms has implications for our understanding of their communication, cooperation, and migration. It has direct functional implications, as controlled cell–cell spacing and ECM continuity will influence antibiotic penetration, phage accessibility, and immune clearance, thereby contributing to the persistence of clinical PA infections.[57]

### 2.3 Cell volume fraction is vertically stratified and not predicted by biofilm thickness

Biofilm thickness or stained biomass is often used as a proxy for biofilm development, but thickness does not reveal how cells are distributed within the matrix.[58–60] To determine whether mucoid and PAO1 biofilms differ in total height and in their depth-resolved cellular organization, we combined cryo-SEM cross-sections with volumetric CLSM analysis of 4-day-old biofilms. Cryo-SEM provided native cross-sectional views including the ECM, whereas CLSM enabled statistical quantification of bacterial volume fractions across the full biofilm depth.

We grew PA biofilms on sapphire discs mounted vertically in the cryo-sample holder and fractured them (**Figure A.8**), thereby allowing a cross-sectional view orthogonal to the growth surface (**Figure A.9)**. This orientation exposes the biofilm profile, from base to surface (**Figure 3a – b**), enabling direct visualization of spatial features such as bacterial layering and changes in density along the biofilm depth, as well as the accurate native biofilm thickness, including ECM. While this approach has been used in cryo-SEM to study biofilm architecture of *Staphylococcus aureus*, *Candida parapsilosis,* and co-cultures of *Staphylococcus epidermidis* and yeast,[21,61] it has not been applied to elucidate the biofilm architecture of PA strains.

**Figure 3:**
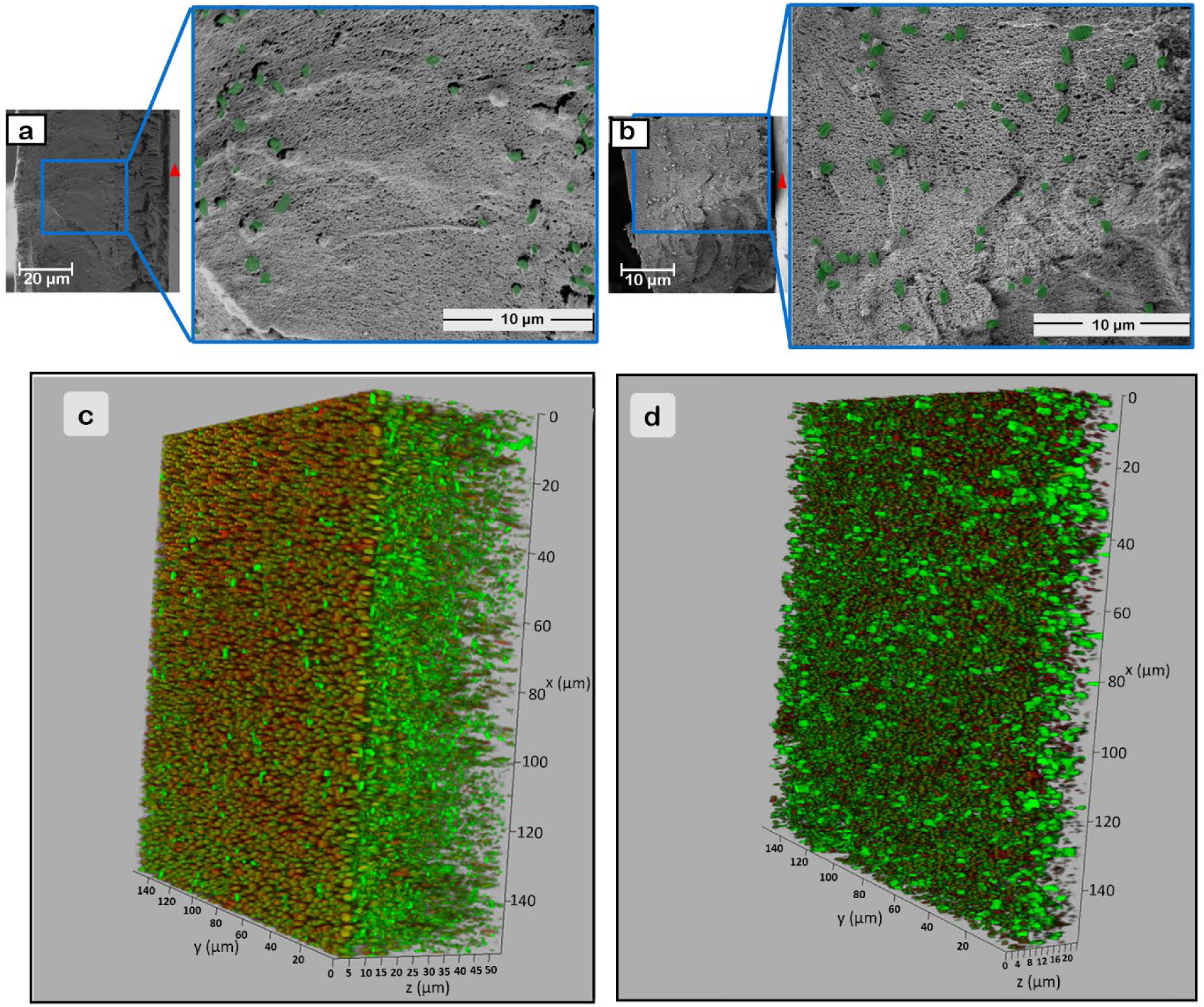
Cryo-SEM micrographs and CLSM images of PA biofilms. PA biofilms were grown on sapphire discs for 4 days and subsequently high-pressure cryo-frozen and fractured orthogonally to the substrate to reveal a biofilm cross-section for a) mucoid PA biofilm and b) PAO1 biofilm. The red triangles point towards the disc surface (substrate). 3D CLSM stacks of c) mucoid PA and d) PAO1 biofilms after 4 days of growth on microscope glass slips.

The cross-sectional cryo-SEM view of biofilms showed that the ECM of both PAO1 and mucoid PA biofilms formed a thick, cohesive, and uniform mesh-like structure (**Figure 3a–b**). The ECM uniformly enveloped the bacterial cells, embedding them within the biofilm. Despite similar matrix appearances, several quantifiable differences in biofilm thickness were observed between the two strains. PAO1 biofilms were thinner, with a mean thickness of 26.8 ± 5.4 µm from the substrate to the top of the ECM after 4 days of growth. Mucoid PA biofilms were thicker (54.5 ± 4.9 µm). CLSM images, in which the thickness was determined by observing fluorescent bacterial cells, yielded similar thicknesses, i.e., 28.27 ± 5.93 µm and 52.88 ± 10.72 µm for PAO1 and mucoid PA, respectively, as shown in **Figure 3c** and **d**. A thickness difference, although less pronounced, was also observed by Nahum *et al*.,[62] who documented 20% higher thicknesses of 5-day-old mucoid PA biofilms (∼65 µm) than non-mucoid PA biofilms.

Quantitative analysis of cell distributions in cryo-SEM cross sections (**Figure A.9**) exhibited substantial variance and inconsistent trends across strains, samples, and regions, particularly in PAO1 biofilms, as it provides only local 2D snapshots of small areas of the biofilm. Quantitative volumetric analysis of CLSM image stacks provided a more robust framework for assessing cell volume fraction across biofilm depth (**Figure 4**). It revealed pronounced spatial variability in cell volume fraction across biofilm regions. While several studies have highlighted elevated bacterial densities and metabolic activities at the biofilm-nutrient source interface,[63–66] to our knowledge, no study has reported the bacterial distribution throughout mucoid PA and PAO1 biofilms.

**Figure 4:**
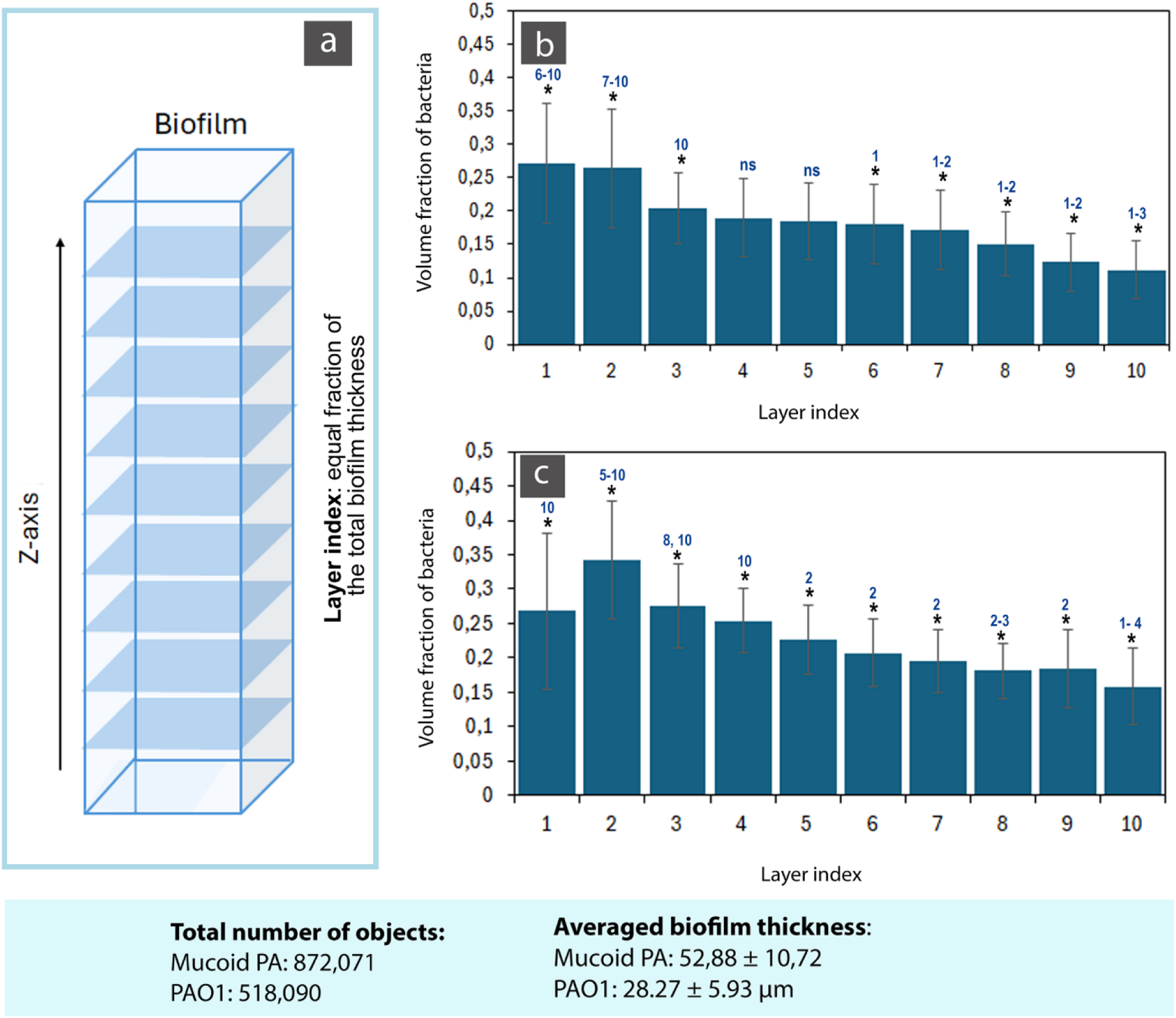
Vertical distribution of bacterial volume fraction across biofilm depth for *P. aeruginosa* biofilms. a) Schematic illustration of the strategy used to generate vertical volume fraction profiles in 3D biofilms. The biofilm volume was partitioned along the z-axis into 10 equally spaced layers, each corresponding to a fraction of the total biofilm thickness. For each layer, the volume fraction of bacteria after segmentation and cylinder fitting was calculated and used to generate averaged vertical density profiles. Bacterial volume fractions per layer for b) mucoid PA and c) PAO1 biofilms are presented as mean ± 95% confidence interval. Statistical comparisons between layers are indicated above the bars (*ns* = not significant (*p* ≥ 0.05); * = significant (*p* ≤ 0.05)).

The total biofilm thickness was divided into 10 equally spaced layers along the z-axis (**Figure 4a**), and for each layer, the bacterial volume fraction was calculated. More than 8 datasets were averaged to generate the mean profiles shown in **Figure 4b–c**. Averaging was performed per layer, ignoring the minor variation in biofilm and layer thickness. The resulting profiles suggest a monotonic decrease in cell volume fraction orthogonal to the surface in both biofilm types. In both cases, the highest bacterial volume fractions were observed in the lower layers (layer indices 1–3), with the lowest-layer fraction (1) significantly higher (*p* ≤ 0.05) than that of the top layer (10), corresponding to an approximately 2-fold reduction in bacterial volume fraction between the most and least densely populated layers. This trend was observed in most individual biofilms analyzed (**Figures A.10 and A.11**), demonstrating that the vertical concentration gradient represents a systematic structural feature rather than an artifact of averaging samples with different gradient profiles and unequal cell numbers.

Most quantitative studies report an increase in bacterial volume fraction toward the bottom/interior of mature PA biofilms, whereas the top is less dense but more metabolically and regulatory active.[32,57,67–69] This stratification is key to understanding persistence and treatment failure. The higher bacterial cell volume fraction at the core of the biofilm is consistent with ongoing cell division, albeit at a reduced rate. The layer-resolved, cell-based occupancy profiles obtained in this study, even for relatively young biofilms, provide quantitative design constraints for developing physiologically relevant 3D biofilm models. Rather than assuming uniform density or homogeneous cell distributions, our data suggest that incorporating non-uniform bacterial concentration profiles into 3D biofilm models is essential for modeling PA biofilms during their formation.

In contrast to our expectations, the average cell volume fractions in the 4-day-old biofilms analyzed with CLSM were similar for PAO1 and mucoid PA, i.e., 0.23 ± 0.06 and 0.19 ± 0.05, respectively (**Figure A.12)**. Although the mucoid PA had a lower volume fraction, the difference was not statistically significant (*p* ≥ 0.05). Because mucoid PA biofilms are significantly thicker than PAO1 biofilms after 4 days but have the same volume fraction, mucoid PA have both undergone greater cell division and produced more ECM. Mucoid PA biofilms have also been reported to have a lower elastic modulus than PAO1,[70] which has been tentatively attributed to a lower cell density. However, our results show a similar cell density, implying that differences in elastic modulus may instead reflect differences in ECM.

Our observations are consistent with prior work showing that PAO1 biofilms have a higher bacterial volume fraction while mucoid *P. aeruginosa* strains produce thicker biofilms and overproduce EPS, particularly alginate.[6,62,71,72,73] However, in our study, this was only indicated and not a statistically significant observation. Henriksen *et al*. also observed that an alginate-overproducing *P. aeruginosa* mucA strain had a lower bacterial volume fraction than the non-mucoid (PAO1) strain.[73] Our weaker differences could indicate that choosing different time points or culture conditions affects such results.

Taken together, the results underscore that biofilm thickness is not a reliable measure of bacterial biomass. Our observation of a higher basal bacterial volume fraction indicates that both biofilm types develop a vertically stratified architecture despite their different total thicknesses. This organization matters because the basal region is expected to have lower nutrient and oxygen availability, stronger matrix confinement, and altered exposure to antimicrobials, whereas the upper region is more directly coupled to the surrounding medium.[10,74,75] It provides valuable quantitative insights into the emergence of biofilm properties, such as robustness, persistence, and treatment tolerance.[10,75] To date, direct, depth-resolved comparisons of biofilm architecture between mucoid PA and PAO1 under comparable experimental conditions have not been reported. Notably, the highest cell volume fraction occurred near the substrate, whereas the strongest clustering was observed near the biofilm surface. Thus, these findings indicate that clustering and cell volume fraction are not equivalent descriptors of biofilm organization, and bacterial cell volume fraction may not automatically predict clustering, although they are often assumed to correlate.

### 2.4 Orientational correlations reveal depth-dependent cell alignment independent of clustering

As PA are asymmetrically shaped and can be represented as oriented cylinders, their local alignment within a 5 µm radius could be quantified using the nematic-order parameter *S*, with *S* = 1 corresponding to perfect nematic order. For close-packed prolate objects, such as PA, orientational ordering has been demonstrated in dense biofilms.[45,46,57,76,77] Mucoid biofilms exhibited weak-to-moderate local orientational order across all three regions, with the nematic-order distributions tending toward higher values near the substrate (**Figure 5a)**. High-order subpopulations reached *S* ≍ 0.85. The overall regional effect for local nematic order in mucoid biofilms showed lower significance after Holm correction (*T*_region_ = 0.0061, adjusted *p* = 0.0552); the increase toward the substrate should therefore be interpreted as a descriptive depth-dependent trend.

**Figure 5:**
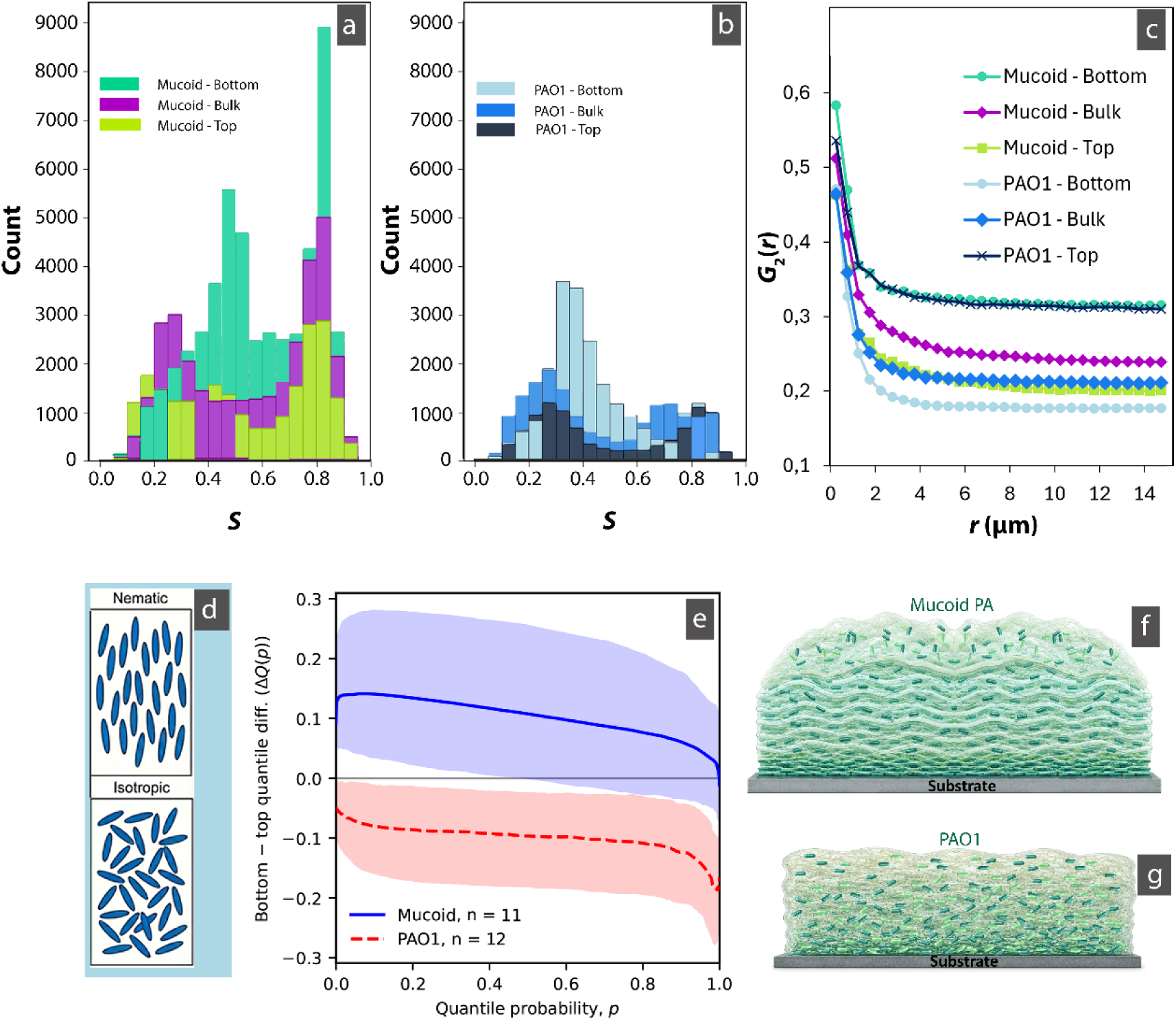
Orientational ordering of 4-day-old mucoid PA and PAO1 biofilms. a – b) Histograms of the local nematic order parameter (*S*). c) The orientational correlation function calculated for the biofilm layers. d) Cartoons for the nematic vs. isotropic order. e) Bottom-to-top mean paired quantile difference across independent 4-day-old mucoid (*n* = 11) and PAO1 (*n* = 9) biofilms. Shaded areas show 95% pointwise bootstrap bands obtained by resampling complete biofilms over 1,000 replicates. Positive values indicate higher nematic order at the bottom, whereas negative values indicate higher order at the top. The reported statistic refers to the omnibus biofilm-type-by-region interaction incorporating bottom, bulk, and top regions. The bottom-to-top curves are shown to illustrate the dominant directional contrast (*T*_interaction_ = 0.0261, Holm-adjusted *p* = 0.0174). f – g) Schematic summarizing the findings regarding mucoid PA and PAO1 biofilm’s depth-dependent cell organization.

Consistent with this descriptive trend, the orientational correlation function, *G*_2_(*r*), decayed more slowly in the bottom layer than in the top layer, indicating that local alignment persisted over longer distances near the substrate (**Figure 5c**). While the cell volume fraction is higher in the lower layers, it is far below the threshold for close-packing to drive orientational ordering and suggests that either the bacterial cells transmit pressure through the ECM or the ordering is the result of coordinated movement of bacteria in preferred local directions that might be a precursor to organized biofilm sub-structures in mature biofilms. Notably, clustering is stronger in the top layers, whereas orientational ordering is stronger in the bottom layers, suggesting that orientational ordering is not simply a consequence of more recent cell division events and residual orientational cell alignment due to a lack of migration. Thus, cell clustering, bacterial volume fraction, and orientational order describe distinct biofilm structural features. Further, the results suggest that early mucoid PA biofilms can develop a coordinated architecture even under sparse, ECM-separated conditions, although image analysis alone cannot reveal the mechanism driving this coordination. These findings extend recent work by Dayton et al. to ECM-separated biofilms without dense cell packing. They demonstrated cellular arrangement in PA14 macrocolony biofilms and showed its effects on substrate distribution, metabolic activity, and antibiotic tolerance.[57]

Although quantitative nematic-order analysis remains limited in *P. aeruginosa* biofilms imaged under similar hydrated conditions,[78] a growing body of biophysical work in other systems demonstrates that orientational organization can emerge independently of simple passive close packing in biofilms.[45–47,77,79] In *Vibrio cholerae*, local liquid-crystalline order and global biofilm morphology have been described to arise from mechanical cell–cell interactions that are modulated by ECM production and external flow.[46] Under confinement, matrix-mediated stress transmission between the biofilm and surrounding material was reported to produce bipolar nematic ordering.[45,76,80,81] Similarly, agent-based modeling predicts that spatially anisotropic stresses near biofilm boundaries can promote cellular alignment, whereas regions dominated by hydrostatic compression are expected to exhibit more disordered packing.[82] Our findings are consistent with observations of long-range nematic ordering of *P. aeruginosa* mediated through the ECM, predominantly close to the substrate.

The depth-dependent distribution of local nematic order (**Figure 5**a–b) differed significantly between mucoid and PAO1 biofilms (*T*_interaction_ = 0.0261, raw permutation *p* = 0.0058, Holm-adjusted *p* = 0.0174; leave-one-biofilm-out sensitivity analysis in **Figure A.13**). The mean bottom-minus-top quantile differences were predominantly positive for mucoid biofilms, indicating a tendency toward greater local order near the substrate, whereas they were predominantly negative for PAO1 biofilms, indicating the opposite depth-dependent profile (**Figure 5**e). PAO1 biofilms also showed a significant overall regional effect (*T*_region_ = 0.0089, Holm-adjusted *p* = 0.0238). No individual pairwise regional contrast remained significant after multiplicity correction. These results show that the two phenotypes differ in their overall spatial profiles of local bacterial alignment, rather than in any particular pair of biofilm regions. The existing literature consistently shows that older PAO1 biofilms are structured, with Psl-rich peripheral regions and internal cavities, whereas alginate overproduction in mucoid strains is associated with thicker, more cohesive, and more persistent biofilm matrices, but these structures occur later, within 4-7 days, and are not directly identifiable with the emergent order imaged here.[83–86]

### 2.5 The mucoid biofilm extracellular matrix exhibits aligned fibrils absent from PAO1 biofilms

Different PA strains do not form the same types of biofilms.[72,87,88] Well-known examples include dense biofilms formed on biomaterial surfaces, where PA acts as an opportunistic pathogen, and mucoid biofilms formed by PA in cystic fibrosis patients.[55,89,90] Despite differences in structure and the critical role of the ECM in shaping biofilm architecture and contributing to antimicrobial resistance, a notable gap remains in our understanding of the ultrastructural organization of different biofilm types. Electron microscopy studies on PA have focused on bacterial cell morphology, surface attachment, or overall biofilm structure.[24,69,91] Differences in internal structure between biofilms, e.g., PAO1 *vs*. mucoid PA, have not been imaged and compared in detail.

Cryo-SEM images from the top, bulk, and bottom regions of mucoid PA revealed structural differences, as shown in **Figure 6**. While the bulk region exhibited mostly locally aligned fibrils extending through the ECM (**Figure 6b**), the top and bottom parts of the biofilm displayed a markedly different ECM morphology. The top and bottom of the biofilm were characterized by a mesh-like appearance of the ECM (**Figure 6a and c**). Although the bottom also displayed fibrils, these were less aligned than in the bulk. We did not find such region-specific differences in ECM architecture among the top, bulk, and bottom zones of mucoid PA biofilms reported in the literature, as these differences are observable only with freeze-fracture cryo-SEM.

**Figure 6:**
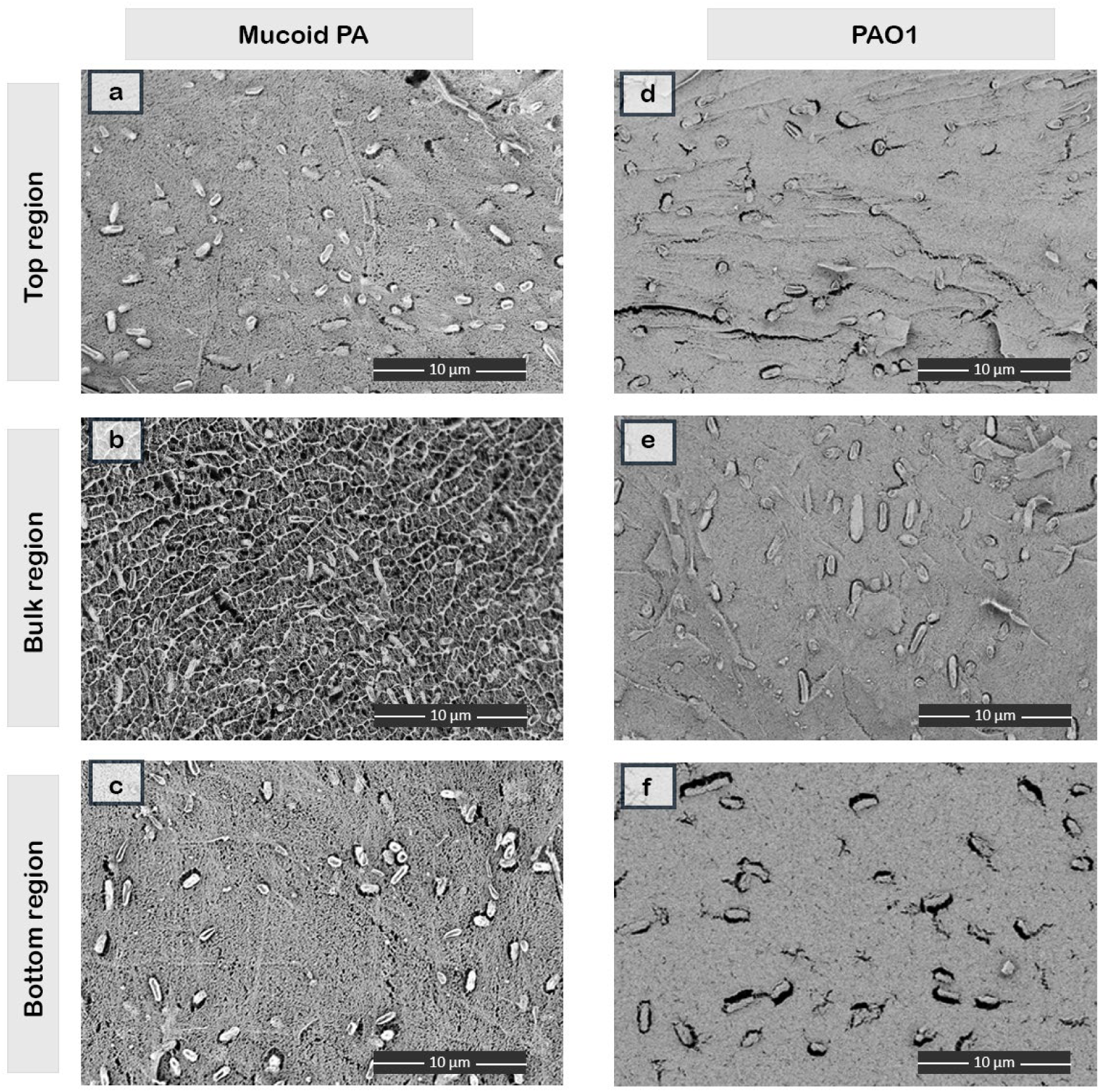
Cryo-SEM micrographs demonstrating variations in ECM architecture in different regions of 4-day-old mucoid and PAO1 biofilms grown on gold carriers. Micrographs shown for a-b) top part of the biofilms (∼1–5 µm from the biofilm surface), c-d) bulk part of the biofilms, and e-f) bottom part of the biofilms (∼1–5 µm from the substrate).

After observing the fibrillar organization in mucoid PA biofilms, we compared these biofilms with PAO1 biofilms to see if such structures were present. The ECM in all three regions of PAO1 biofilms presents a dense and mesh-like structure (**Figure 6d – f)**, suggesting that the aligned fibrillar organization is typical for the mucoid strain but not ubiquitously observed.

The observations of aligned fibrillar structures in the interior of mucoid PA biofilms but not in the denser PAO1 biofilms were surprising, as the ECM in mucoid biofilms is distinguished by predominantly being formed from alginate, which has more than an order of magnitude smaller dimensions than the thickness and spacing of the structures observed in our images. Notably, the aligned fibrillar features are also absent in compositionally comparable alginate hydrogels (**Figure A.14**), suggesting that these features observed in mucoid PA biofilms do not arise from the alginate matrix but instead reflect a biofilm–specific structural organization involving other ECM components that self–assemble or are organized by the mucoid PA during matrix synthesis. While the cryo–SEM data do not allow direct assignment of biochemical identity, the contrast, size, and organization of these fibrillar structures are consistent with larger molecular assemblies, such as bundles formed by large glycoconjugates, eDNA complexes, amyloid-forming proteins (e.g., Fap fibers), or glycosylated adhesins, which are known ECM components of *Pseudomonas*.[92–94] These components are known to self-assemble into filamentous or fibrillar structures and could possibly contribute to the observed morphology.[95–99] We note that some well-characterized fibrillar matrix proteins, such as CdrA, are primarily associated with Psl-rich, non-mucoid biofilms, such as PAO1, and may play a reduced or altered role in alginate-dominated mucoid strains.[72,100] Nevertheless, strain-specific expression patterns do not fully exclude their involvement, particularly in mature biofilms, where multiple matrix components may coexist and interact.

Filamentous matrix molecules are increasingly recognized as key determinants of biofilm viscoelasticity and cohesion; our observation of their enrichment in the bulk suggests they are part of the load–bearing scaffold that allows mucoid biofilms to resist shear and deformation.[96,101] Conversely, aligned fibers might lead to weak shear planes, promoting shedding of biofilm sheets under shear stress. Furthermore, the formation of fibrous structures may be linked to biofilm maturation, increased matrix complexity, or mechanical confinement, which may promote higher-order assembly.[102,103] Ultimately, the mature, dense, cohesive volumes of biofilms are where nutrient gradients, slow growth, and the highest antibiotic tolerance reside,[104,105] linking matrix ultrastructure to the formation of highly protected “refuge” compartments for cells. Previous studies have shown that disruption of matrix architecture through enzymatic degradation of extracellular DNA, inhibition of amyloid fiber formation, or interference with polysaccharide synthesis can significantly enhance antibiotic efficacy and immune clearance.[106–108]

Definitive identification of the molecular composition of observed features will require complementary biochemical, proteomic, or targeted labeling approaches, which are beyond the scope of the present study but represent important directions for future work.

## 3 Conclusion

Our results revise the structural view of young, 4-day-old *P. aeruginosa* biofilms. Rather than forming densely packed microcolonies, both mucoid and PAO1 biofilms were dominated by spatially separated cells embedded in continuous ECM. Native-state cryo-SEM preserved this architecture, whereas dehydration-based preparation introduced matrix collapse and apparent cell clustering, underscoring the risk of artifacts in conventional SEM.

Quantitative cryo-SEM and CLSM analyses showed that biofilm organization varies with depth and cannot be described by a single structural metric. Bacterial volume fraction was highest near the substrate, whereas clustering was stronger toward the surface. Orientational analysis further indicated depth-dependent cell alignment that was not simply explained by either cell density or clustering.

Together, these results show that early PA biofilms are vertically stratified, ECM-separated cellular systems rather than homogeneous aggregates. The mechanisms underlying this stratification remain unresolved, but the observed architecture is consistent with depth-dependent growth, matrix constraints, and matrix-mediated, local cell–cell interactions.

Mucoid PA and PAO1 differed in matrix architecture. Mucoid biofilms contained aligned fibrillar ECM structures in the biofilm interior, whereas PAO1 biofilms exhibited a denser mesh-like matrix. Although the molecular identity of the fibrillar structures remains unknown, their strain-specific occurrence suggests higher-order matrix organization beyond alginate alone. While mucoid PA biofilms were twice as thick as PAO1 biofilms and more stratified, their cell volume fractions were similar.

Our findings establish native-state cryo-SEM combined with volumetric CLSM and spatial statistics as a quantitative framework for biofilm structural biology. They provide a corrected structural basis for interpreting biofilm organization, tolerance, transport, phage and immune access, cell–cell interactions, and the design of more realistic experimental and computational biofilm models, although generalization will require testing additional biofilm ages and growth conditions.

## 4. Materials and Methods

### Bacterial strains, growth conditions, and antibiotics

The *P. aeruginosa* (PA) strains utilized in this study include PAO1(ATCC 15692) and a mucoid strain (ATCC 39324). Unless otherwise specified, frozen bacterial stocks were thawed and initially cultured overnight at 37 °C in Tryptic Soy Broth (TSB). Subsequently, the bacteria were sub-cultured onto Tryptic Soy Agar (TSA) plates and incubated for 24 h. Colonies were then transferred to 10 ml of TSB and cultured at 37 °C for 16–18 h with agitation at 100 rpm to produce a bacterial suspension for further experimentation.

### Biofilm growth

Overnight cultures of PA strains were harvested and washed 3 times with PBS. The bacterial suspensions were then adjusted to an optical density at 600 nm (*OD*₆₀₀) of 0.1 in TSB. Sterile 18 x 18 mm cover glasses (thickness 1 – Carl Roth), 3 mm diameter specimen carriers (gold-coated – Type A, Leica Microsystems Inc., Austria) with 100 µm indentations, along with 6 mm sapphire discs (Leica Microsystems Inc., Austria), were placed in Petri dishes containing the respective bacterial suspensions. Unless otherwise stated, PA strains were allowed to grow for 4 days in TSB to form biofilms on these substrates at 37 °C.

Developed biofilms on Type A carriers for SEM/cryoSEM were further covered with an additional 3 mm, carefully lined with double-sided tape to create a secure sandwich after 4-day growth prior to sample preparation. The sapphire discs with biofilms were placed (biofilm side up) into the indentation of a 6 mm Type A carrier. A spacer was added on top to prevent movement and ensure even contact. The assembly is completed by placing a matching top carrier over the disc, creating a secure sandwich.

### Sample preparation for CLSM and SEM

*P. aeruginosa* biofilms (on cover glasses) were prepared for confocal laser scanning microscopy (CLSM) following incubation under the conditions described above. To preserve biofilm structure while enabling fluorescence imaging, samples were gently handled throughout preparation.

Biofilms were stained using a Live/Dead BacLight bacterial viability kit (Thermo Fisher Scientific) according to the manufacturer’s instructions. Following staining, rinsing of PA biofilms grown on cover glasses was avoided to prevent shear-induced disruption and loss of loosely attached cells; instead, excess staining solution was carefully removed from the staining chamber by pipetting. The biofilm samples were afterward overlaid with an additional coverslip to prevent evaporation during CLSM acquisition. Imaging was performed using a Leica TCS SP8-STED inverted microscope.

For SEM, PA biofilms were prepared by AD, CPD, or HPF (see appendixes for a more detailed description).

## Acknowledgment and Funding

This project was supported by the BOKU Core Facility Multiscale Imaging. We thank the FWF Austrian Science Fund for funding (P33226).

## Code Availability

The underlying custom Python code for this study is not yet publicly available but may be made available upon reasonable request to the corresponding author.

