## Supplementary information for "Native-State Imaging Reveals Spatially Separated Organized Cells and Strain-Specific Matrix Architecture in *Pseudomonas aeruginosa* Biofilms"

Running Head: Cell and matrix organization in *P. aeruginosa biofilms*

[#] Present address: Schandl Stephan, Department of Chemistry and Biology, School of Science and Technology, University of Siegen, Siegen Germany

[#] Present address: Olivier Guillaume, Pre-award Support Department, BOKU University, Vienna, Austria.

### APPENDIX

#### Materials and Methods

##### Sample preparation for SEM imaging

*Air/HMDS drying and scanning electron microscope imaging of samples*

After incubation, PA biofilms were chemically fixed by immersion in a 2.5% glutaraldehyde solution (Sigma) and left overnight at 4 °C. Following fixation, the samples underwent two consecutive rinses with a 0.9 wt% sodium chloride (NaCl) solution, followed by sequential dehydration in ethanol solutions of increasing concentrations: 30%, 50%, 70%, 90%, 95%, and finally, 100%, all conducted at room temperature for 3 min each. Subsequently, the chemically fixed samples were treated with 100% hexamethyldisilazane (HMDS) for 1 min and air-dried. Prior to air-drying, biofilms grown on sample carriers were manually fractured using a clean blade to expose internal structures. Similarly, biofilms grown on glass substrates were carefully cut into smaller sections following air-drying to facilitate subsequent imaging. All samples were uniformly sputter-coated (Leica EM SCD005, Leica, Germany) with a ~8 nm thick layer of gold to enhance sample conductivity and image contrast. SEM imaging was carried out using an Apreo VS SEM (Thermo Scientific, The Netherlands) at an acceleration voltage of 5.0 kV and a current of 0.10 nA. High vacuum back-scattered and secondary electrons were detected using T1 and T2 in-lens detectors, respectively.

*Critical point drying and scanning electron microscope imaging of samples*

PA biofilms were subjected to chemical fixation and dehydration as described above. Subsequently, samples were transitioned to acetone (Sigma) as the intermediate solvent before drying using a critical-point dryer (Leica EM CPD030, Leica, Germany). Biofilms grown on sample carriers were manually fractured with a clean blade after CPD to expose internal structures. Biofilms grown on glass substrates were carefully cut into smaller sections before CPD to accommodate the dimensions of the drying chamber. All samples were subsequently coated with ~8 nm of gold and then imaged using an SEM (Apreo VS, Thermo Scientific, The Netherlands).

*Cryo preparation and scanning electron microscope imaging of samples*

All sandwiched carriers were immediately loaded into the Leica EM HPM100 high-pressure freezer (Leica Microsystems Inc., Austria) and automatically frozen according to the protocol described in Garbarino et al.[1] The frozen samples were subsequently placed in a liquid-nitrogen-filled Dewar for preservation before being transferred to the freeze-fracture system under cryogenic conditions. The frozen sample carriers or sapphire discs were mounted onto the sample holder under liquid nitrogen ($LN_2$) and transferred to a Leica EM ACE900 freeze-fracture system (Leica Microsystems Inc., Austria). Freeze-fracturing was performed at -120 °C using a flat-edge knife.

Unless otherwise stated, to enhance structural visibility of fractured samples, sublimation (etching) was performed for 20 s at –110 °C under a vacuum of ~$5.6 \times 10^{-7}$ mbar. Following the sublimation step, the fractured samples were e-beam coated with a 4-nm platinum layer at a 45° angle and a 4-nm carbon layer at a 90° angle to improve contrast and minimize beam damage during imaging. Subsequently, the samples were transferred to SEM via a cryo-transfer system (Leica EM VCT500).

Cryo-SEM imaging was performed using the Apreo VS SEM (Thermo Scientific, The Netherlands) at an operating voltage of 1.0 kV and a current of 0.10 nA in a high vacuum mode. Backscattered and secondary electrons were detected via the in-lens T1 and T2 detectors, respectively. Throughout the imaging process, the SEM chamber temperature was maintained between -110 °C and -120 °C, depending on the chamber pressure to avoid sublimation and condensation, as observed *in situ*.

For each imaging technique, more than three independent samples were prepared. From each sample, a minimum of 10 fields of view were imaged, and representative images were selected based on the preservation of sample structures. Where applicable, bacterial cells were manually false-coloured green using Adobe Photoshop (version 25.5.3).

#### Quantitative spatial analysis of *Pseudomonas* biofilm structure

##### *Cryo-SEM data processing*

Cryo-SEM images were analyzed consistently using a custom Python script applied to the overlay images derived from the cryo-SEM datasets. Individual bacterial cells were first manually masked (**Figure A1**), as the dense and heterogeneous biofilm architecture precluded reliable automated cell detection using standard image analysis software (e.g., ImageJ) or commonly used segmentation algorithms. The manually masked images served as the input for subsequent computational analysis.

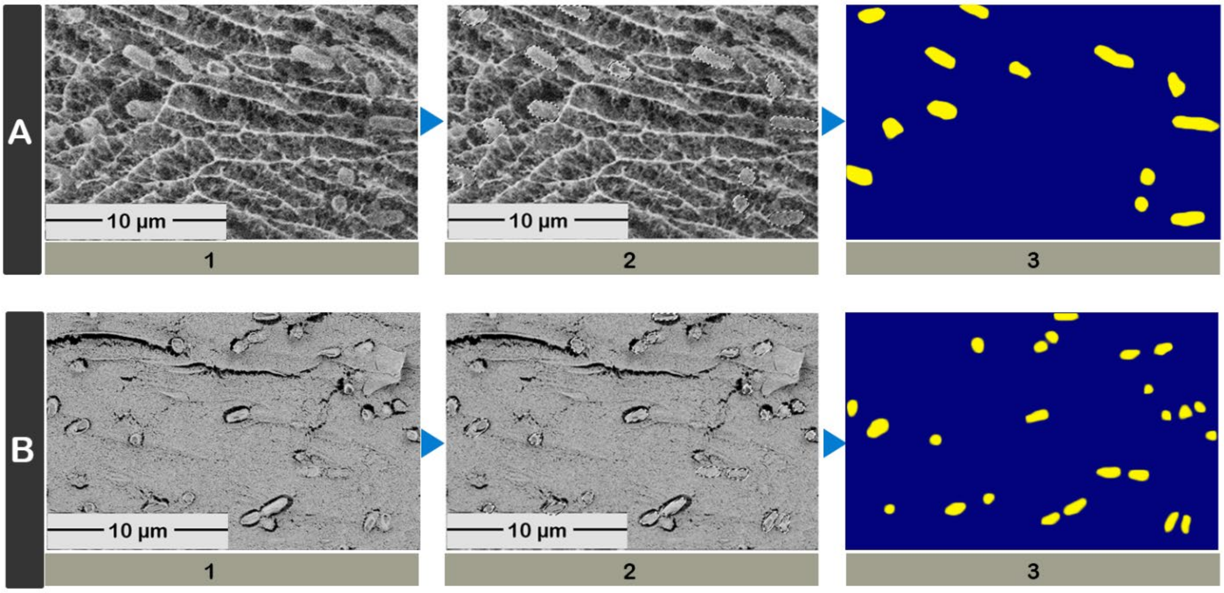


**Figure A1**: Workflow for cryoSEM image processing and extraction of bacteria for spatial analysis. (1) Representative cryo-SEM images of PA biofilms (A and B) showing native ECM architecture and embedded bacterial cells. (2) Manually masked bacterial cells. (3) Masked bacterial cells following color thresholding.

To isolate bacterial shapes, the Python script applied a color-based threshold to identify the yellow hue corresponding to the masked bacteria. The resulting pixel binary masks underwent a morphological opening (dilation followed by erosion) to remove small artifacts, suppress noise, and smooth object boundaries prior to spatial analysis. Geometric features were then extracted by identifying contours within the processed binary masks. For each contour, the script fitted an ellipse to determine the centroid coordinates, major-axis length, minor-axis thickness, and orientation angle of each bacterium. The extracted centroid coordinates were subsequently used for spatial analyses.

*CLSM data processing*

CLSM data were analyzed consistently, without manual steps, using a custom Python script applied to the raw voxel fluorescence intensity data from each biofilm measurement. In brief, an intensity threshold was applied to the combined fluorescence intensity of the live and dead channels to isolate bacterial signals from background noise for each voxel. Bacteria were identified by segmenting the merged fluorescence signals using a Watershed algorithm. A cylindrical-shape constraint was used, requiring a cylindrical shape with maximum and minimum lengths and radial dimensions, defined by compiling histograms of bacterial dimensions for PAO1 and mucoid PA, respectively, from cryo-SEM data for each strain and adding the microscopy point-spread functions in x, y, and z, respectively. The cylinders were fitted using weighted principal component analysis, accounting for the intensity of individual voxels rather than a simple binary mask. Bacteria with overlapping fluorescence signals were separated by recursively splitting objects, rejecting too thin objects, and forcing splitting by halving the length of objects if they are outside of the shape constraints after reaching the maximum depth of the Watershed algorithmic splitting. The resulting data on bacteria annotated with cylinders were overlaid on the raw fluorescence data for classification based on their fractions of live and dead fluorescence, and their spatial distributions were used to calculate centroids and volumes for statistical analysis of CLSM data.

*Radial distribution function (RDF) and nearest-neighbor distance analyses*

To assess cell–cell spacing in PA biofilms, the $RDF$ (with Monte Carlo correction – **equation 1**), defined as the normalized probability density of finding a neighbor at distance $r$ from a reference particle, compared to a random distribution, was calculated from data obtained from the processed cryo-SEM (2D radial coordinates) and CLSM (cylindrical coordinates) images, respectively.

$$g(r) = \frac{H_{obs}(r)}{H_{rand}(r)} \times \frac{N_{rand}}{N_{obs}} \quad (1)$$

Python scripts calculated the nearest-neighbor distances, defined as the Euclidean distance from each bacterium's centroid to that of its closest neighbor in 2D for cryo-SEM data and in 3D for CLSM data.

$RDF$s from datasets of the same sample type (strain and imaging technique) were aggregated, and the mean $g(r)$ was calculated together with the 95% confidence intervals, assuming a Student's t-distribution. Nearest-neighbor distances were summarized as histograms, from which mean and modal (peak) values were extracted.

*Vertical cell density profiling*

To evaluate how cell density varied with distance from the substrate, cryo-SEM images of fractured biofilms grown on sapphire discs were analyzed. Each image was divided into horizontal rectangular regions of interest (ROIs), aligned parallel to the substrate. For mucoid biofilms, ROI widths were set to 5 µm, while for PAO1 biofilms, a width of 4 µm was used to accommodate differences in biofilm structure and cell density. Cell counts within each ROI were manually determined in Adobe Photoshop to ensure consistent and accurate quantification. The constant area across ROIs enabled direct calculation of cell density (cells per μm$^2$) as a function of distance from the substrate.

Additionally, vertical cell-density profiles were derived from 3D CLSM image stacks to quantify the spatial distribution of bacterial cells along the biofilm thickness. Within the entire analyzed volume, bacterial cells (fitted cylinders) were identified using intensity-based segmentation described above. The total biofilm thickness was divided into 10 equally spaced layers along the z-axis, and for each layer, the volume fraction of cylinders fitted to represent bacteria was calculated. Multiple datasets (>8) from both PA strains were subsequently averaged to generate the mean profiles. Averaging was performed layer by layer, ignoring the minimal variation in biofilm and layer thickness. Layer-resolved volume fractions were aligned by layer index, as all datasets were discretized into the same number of Z layers.

For statistical evaluation, mean density and 95% Confidence Interval (CI) were calculated for each bin. One-way ANOVA with Tukey and LSD post hoc tests was performed to determine significant differences in cell density across depths. (IBM SPSS Statistics (27.0.1)). A $p$-value ≤0.05 was considered statistically significant.

*Ripley's H(r) spatial analysis*

Clustering of bacteria in biofilm was quantified using Ripley's $H(r)$ function (**equation 2**).

$$K(r) = \int_0^r g(r')4\pi(r')^2dr' \; ; \; L(r) = \left(\frac{K(r)}{\frac{4}{3}\pi}\right)^{1/3} \; ; \; H(r) = L(r) - r \qquad (2)$$

To assess depth-dependent organization, biofilms were subdivided into three volume regions based on their vertical position: the biofilm–substrate interface (Bottom), the biofilm interior (Bulk), and the biofilm–liquid interface (Top). Each was designated as 20% of the biofilm thickness in the corresponding region: 0-20% (Bottom), 40-60% (Bulk), and 80-100% (Top). Ripley's $H - function$ was calculated for each region. $H(r) \approx 0$ indicates random spatial organization, positive $H(r)$ values indicate clustering, negative $H(r)$ values indicate spatial inhibition or regularity, and the peak in $H$ gives an approximate cluster radius.

To ensure robust comparison between strains, Ripley's $H(r)$ functions were averaged across at least six independent biological replicates per strain. All analyses were performed using Python scripts on the centroid positions of bacteria identified in the 3D confocal data.

*Nematic Order Parameter* ($S$)

Bacterial orientation was represented by the unit vector along the major axis of each fitted cylinder, $\boldsymbol{u}_i$, determined by principal component analysis. To quantify bacterial alignment in the biofilm, we used the Q-tensor approach from liquid crystal physics (**equation 3)**. This robustly handles the fact that a bacterium pointing "up" is equivalent to one pointing "down" (head-tail symmetry).

$$\boldsymbol{Q} = \frac{1}{N}\sum_{i=1}^{N}\left(\frac{3}{2}\boldsymbol{u}_i \times \boldsymbol{u}_i - \frac{1}{2}\mathbf{I}\right) \tag{3}$$

Diagonalizing $Q$ yields $S$ (the scalar nematic order parameter), which is the largest eigenvalue. It ranges from 0 (isotropic/random) to 1 (perfect alignment). Local nematic order parameters were calculated for bacteria with at least three nearest neighbors within a 5 µm radius; regional nematic order parameters were calculated for the entire regional volume (bottom, bulk, and top), and the nematic order parameter of the biofilm was calculated from $Q$ yields $S$ for all bacteria in the biofilm.

Local nematic-order distributions were analyzed using independently grown biofilms as the unit of inference ($n = 11$ mucoid and $n = 9$ PAO1 biofilms). For each biofilm and each vertical region—bottom, bulk, and top—all available local $S$ values were represented by an empirical quantile function evaluated on a common non-endpoint grid of 10,000 probabilities, $p_k = (k + 0.5)/10{,}000$. Each biofilm contributed one quantile function with equal weight, irrespective of the number of local measurements. Overall regional effects within each phenotype were tested using a permutation statistic, $T_{\mathrm{region}}$, based on the integrated squared deviations of the three regional mean quantile functions, with region labels permuted only within individual biofilms. The biofilm-type-by-region interaction was tested using $T_{\mathrm{interaction}}$, calculated from differences between the centered regional mean quantile functions of the two phenotypes; complete three-region biofilm profiles were reassigned between phenotypes over 10,000 permutations. Pairwise regional comparisons within each phenotype used exact sign-flip tests of the complete paired quantile-difference functions, whereas phenotype comparisons used sample-level label permutations. Pairwise distributional effect sizes were reported as 2-Wasserstein distances ($W_2$), with 1-Wasserstein distances ($W_1$) included as descriptive robustness measures. Holm correction was applied separately to the three nematic-order omnibus tests, the six within-phenotype regional comparisons, and the four between-phenotype comparisons. For visualization, mean bottom-minus-top quantile-difference curves and 95% pointwise bootstrap bands were calculated by resampling complete biological biofilms within each phenotype over 1,000 bootstrap replicates. The bootstrap bands were used descriptively and not for pointwise hypothesis testing.

Bacterial orientation was represented by the unit vector along the major axis of each fitted cylinder, $\boldsymbol{u}_i$, determined by principal component analysis. Orientational alignment was quantified using the nematic $Q$-tensor formalism from liquid-crystal physics.[2] Local nematic order parameters were calculated for a local neighborhood $\mathcal{N}_i$ for each bacterium with at least three nearest neighbours within a 5 μm radius; regional nematic order parameters were calculated for the entire regional volume (bottom, bulk, and top), and the nematic order parameter of the biofilm was calculated from $Q$ yields $S$ for all bacteria in the biofilm. The local $Q$-tensor was calculated as:

$$\mathbf{Q}_i = \frac{1}{|\mathcal{N}_i|}\sum_{j\in\mathcal{N}_i}\left(\frac{3}{2}\mathbf{u}_j\mathbf{u}_j^T - \frac{1}{2}\mathbf{I}\right), \quad (4)$$

where $\mathbf{I}$ is the $3 \times 3$ identity matrix. The local nematic-order parameter $S_i$ was defined as the largest eigenvalue of $\mathbf{Q}_i$. Values close to 0 indicate locally isotropic orientations, whereas values close to 1 indicate strong local alignment.

For depth-resolved analysis, each biofilm was divided into three vertical regions: bottom, bulk, and top, corresponding to the biofilm–substrate interface (0-20% of the biofilm thickness), biofilm interior (40-60% of the biofilm thickness), and biofilm–liquid interface (80-100% of the biofilm thickness), respectively. For each independently grown biofilm and each region, all available local $S_i$values were retained and analyzed as an empirical distribution. The biological biofilm ($n = 11$ mucoid and $n = 9$ PAO1 biofilms), not the individual local measurement, was used as the unit of statistical inference.

To compare complete distributions rather than only means or peak positions, each sample–region distribution was represented by its empirical quantile function, $Q_{igr}(p)$, where $i$ denotes biological sample, $g$ phenotype, $r$ region, and $p$ quantile probability. Quantile functions were evaluated on a common non-endpoint grid of 10,000 probabilities:

$$p_k = \frac{k+0.5}{10{,}000}, k = 0, \dots, 9999. \quad (5)$$

Each biological sample contributed one quantile function per region and was weighted equally in group-level calculations, irrespective of the number of local measurements. Group-level distributions were summarized by one-dimensional Wasserstein barycentres, obtained by averaging the sample-specific quantile functions. Pairwise distributional effect sizes were reported as 2-Wasserstein distances, $W_2$, which in one dimension can be calculated directly from empirical quantile functions[3]:

$$W_2(A, B) = \left[\frac{1}{K} \sum_{k=1}^{K} (Q_A(p_k) - Q_B(p_k))^2\right]^{1/2}. \quad (6)$$

The 1-Wasserstein distance:

$$W_1(A,B) = \frac{1}{K}\sum_{k=1}^{K} | Q_A(p_k) - Q_B(p_k) |, \quad (7)$$

was calculated as a descriptive robustness measure. The sample-level regional quantile functions and corresponding Wasserstein barycentres are shown in **Figure A2**.

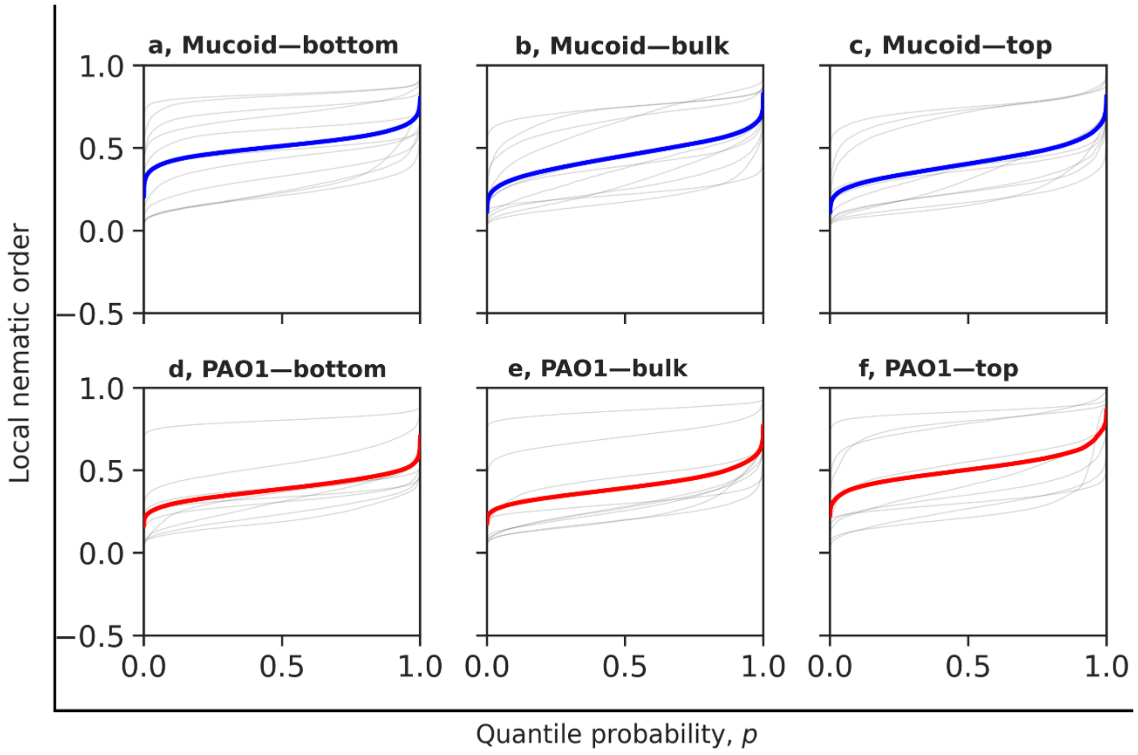


**Figure A2:** Regional quantile functions and Wasserstein barycentres for local nematic order. Thin lines show empirical quantile functions for individual biological biofilms in the bottom, bulk, and top regions. Thick lines show equally weighted Wasserstein barycentres for each phenotype and region. Each biological biofilm contributes one quantile function with equal weight, irrespective of the number of local nematic-order measurements

Regional effects within each phenotype were tested using restricted permutation tests.[4] For each phenotype, the omnibus regional test statistic was calculated from the integrated squared deviations of the three regional mean quantile functions from their phenotype-specific mean profile:

$$T_{\text{region}} = \sum_r \int \left[\bar{Q}_{gr}(p) - \bar{Q}_{g\cdot}(p)\right]^2 dp. \quad (8)$$

The null distribution was generated by permuting the bottom, bulk, and top labels only within each biological sample, preserving the matched regional structure of each biofilm. $T_{\text{region}}$ measures how strongly the three regional distributional barycentres differ within one phenotype. A value of $T_{\text{region}} = 0$ indicates that the bottom, bulk, and top barycentre quantile functions are identical. Larger values indicate larger distributional separation between regions, with larger discrepancies weighted more strongly because the statistic is based on squared differences. The statistic is expressed in squared units of the analyzed order parameter and is used as an omnibus test statistic, not as a pairwise Wasserstein distance. For an order parameter bounded to the interval $[a, b]$, with range $R = b - a$, $T_{\text{region}}$ is bounded by:

$$0 \leq T_{\text{region}} \leq \frac{2}{3}R^2. \quad (9)$$

For the local nematic-order parameter $S \in [0,1]$, this corresponds to:

$$0 \leq T_{\text{region}} \leq \frac{2}{3}. \quad (10)$$

The biofilm-type-by-region interaction was tested using an omnibus statistic calculated from the difference between the centered regional profiles of the two phenotypes:

$$T_{\text{interaction}} = \sum_r \int \left[\left(\bar{Q}_{\text{mucoid},r}(p) - \bar{Q}_{\text{mucoid},\cdot}(p)\right) - \left(\bar{Q}_{\text{PAO1},r}(p) - \bar{Q}_{\text{PAO1},\cdot}(p)\right)\right]^2 dp. \quad (11)$$

For this interaction test, complete three-region biofilm profiles were randomly reassigned between phenotypes over 10,000 permutations. Individual biological-sample bottom-minus-top quantile-difference curves are shown in **Figure A3**.

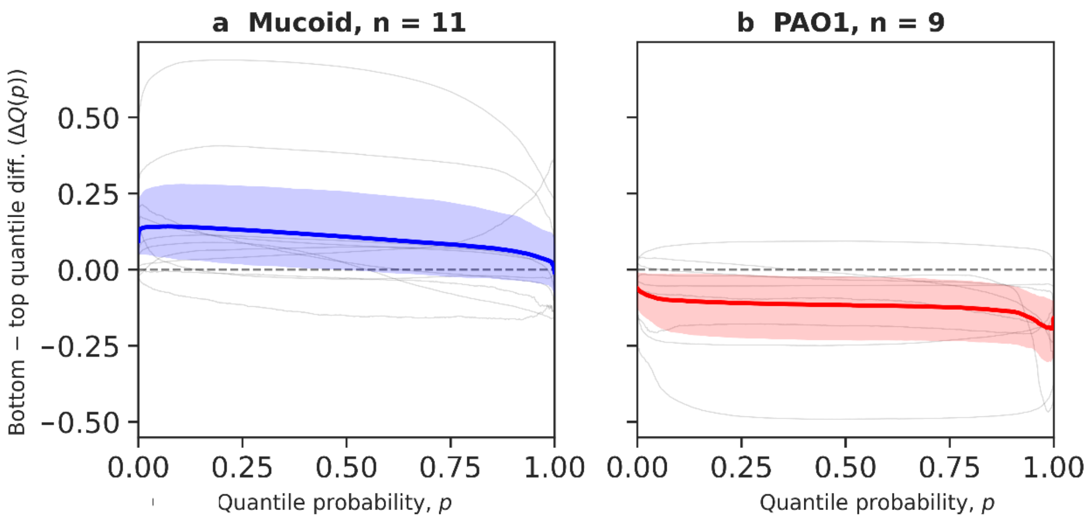


**Figure A3:** Individual bottom-to-top quantile-difference curves for local nematic order. Thin lines show the paired quantile-difference curves, ΔQ_i (p)=Q_(i,bottom) (p)-Q_(i,top) (p), for individual mucoid and PAO1 biofilms. Thick lines show phenotype-specific mean paired differences, and shaded regions show 95% pointwise bootstrap bands obtained by resampling complete biological biofilms. Positive values indicate higher local nematic order at the bottom of the biofilm.

This statistic tests whether the depth-dependent regional profile of nematic-order distributions differs between mucoid and PAO1 biofilms after removing any phenotype-wide shift in nematic order. A value of $T_{\mathrm{interaction}} = 0$ indicates that the centered bottom–bulk–top distributional profiles are identical between phenotypes, even if the two phenotypes differ by an overall shift. Larger values indicate stronger differences in the regional pattern. For an order parameter bounded to $[a, b]$:

$$0 \leq T_{\mathrm{interaction}} \leq \frac{8}{3} R^2, \tag{12}$$

and for $S \in [0,1]$:

$$0 \leq T_{\mathrm{interaction}} \leq \frac{8}{3}. \tag{13}$$

Because both $T_{\mathrm{region}}$ and $T_{\mathrm{interaction}}$ are squared omnibus statistics, their absolute numerical values should be interpreted relative to the order-parameter scale and the permutation null distribution. For descriptive interpretation, $\sqrt{T/3}$ gives an approximate root-mean-square quantile-profile deviation per region on the original $S$-scale. Statistical significance was determined only from the corresponding restricted permutation tests. All integrals were evaluated numerically as averages over the common 10,000-point quantile grid.

Post hoc regional comparisons within each phenotype were performed on complete paired quantile-difference functions:

$$D_i(p) = Q_{ir_1}(p) - Q_{ir_2}(p), \quad (14)$$

using exact sign-flip permutation tests. All sign configurations were enumerated for the present sample sizes. Region-specific and complete-biofilm phenotype comparisons were tested using biological-sample-level label permutations. Family-wise error rates were controlled using Holm's sequential procedure.[5] Holm correction was applied separately to the three nematic-order omnibus tests, the six within-phenotype regional post hoc comparisons, and the four between-phenotype comparisons.Complete statistical results, including omnibus tests, pairwise Wasserstein distances, raw $p$-values, Holm-adjusted $p$-values, and permutation procedures, are provided in **Table A1**.

For visualization, mean bottom-minus-top quantile-difference curves:

$$\Delta Q(p) = Q_{\text{bottom}}(p) - Q_{\text{top}}(p), \quad (15)$$

were plotted for mucoid and PAO1 biofilms. Positive values therefore indicate higher nematic order at the bottom, whereas negative values indicate higher nematic order at the top. Descriptive 95% pointwise bootstrap bands were generated by resampling complete biological biofilms with replacement within each phenotype over 1,000 bootstrap replicates. These bootstrap bands were used only to visualize between-biofilm variability and were not used as pointwise hypothesis tests. All analyses were performed using a custom Python script.

**Table A1. Complete Wasserstein-based statistical analysis of local nematic-order distributions.** The table reports omnibus regional and biofilm-type-by-region tests, pairwise regional comparisons, phenotype comparisons, $T_{\mathrm{region}}$, $T_{\mathrm{interaction}}$, $W_2$, descriptive $W_1$, raw $p$-values, Holm-adjusted $p$-values, sample sizes, and permutation procedures.

| Analysis_Family | Comparison | Metric | n_Mucoid | n_PAO1 | n_Pairs | Effect_Size_Type | Effect_Size | W1_Descriptive | Descriptive_Bootstrap Range | Raw_p | Holm_Adjusted_p | Test_Type | Number_of_Permutations |
|---|---|---|---|---|---|---|---|---|---|---|---|---|---|
| Omnibus (Primary) | Mucoid (Bot=Blk=Top) | nematic_order | 11 | - | - | $T_{\mathrm{region}}$ | 0,006 | | - | 0,055 | 0,055 | Monte Carlo region label permutation | 10000 |
| Omnibus (Primary) | PAO1 (Bot=Blk=Top) | nematic_order | - | 9 | - | $T_{\mathrm{region}}$ | 0,009 | | - | 0,012 | 0,024 | Monte Carlo region label permutation | 10000 |
| Omnibus (Primary) | Biofilm x Region Interaction | nematic_order | 11 | 9 | - | $T_{\mathrm{interaction}}$ | 0,026 | | - | 0,006 | 0,017 | Monte Carlo complete profile permutation | 10000 |
| PostHoc Intra (Mucoid) | Bottom vs Bulk | nematic_order | 11 | - | 11 | W2 | 0,070 | 0,062 | (0.0270, 0.1409) | 0,079 | 0,316 | Exact paired sign-flip | 2048 |
| PostHoc Intra (Mucoid) | Bottom vs Top | nematic_order | 11 | - | 11 | W2 | 0,108 | 0,103 | (0.0273, 0.2309) | 0,098 | 0,316 | Exact paired sign-flip | 2048 |
| PostHoc Intra (Mucoid) | Bulk vs Top | nematic_order | 11 | - | 11 | W2 | 0,042 | 0,041 | (0.0041, 0.1298) | 0,467 | 0,934 | Exact paired sign-flip | 2048 |
| PostHoc Intra (PAO1) | Bottom vs Bulk | nematic_order | - | 9 | 9 | W2 | 0,019 | 0,013 | (0.0113, 0.0692) | 0,594 | 0,934 | Exact paired sign-flip | 512 |
| PostHoc Intra (PAO1) | Bottom vs Top | nematic_order | - | 9 | 9 | W2 | 0,121 | 0,120 | (0.0403, 0.2376) | 0,035 | 0,176 | Exact paired sign-flip | 512 |
| PostHoc Intra (PAO1) | Bulk vs Top | nematic_order | - | 9 | 9 | W2 | 0,107 | 0,107 | (0.0323, 0.2269) | 0,023 | 0,141 | Exact paired sign-flip | 512 |
| PostHoc Whole-Biofilm | Mucoid vs PAO1 (Complete) | nematic_order | 11 | 9 | - | W2 | 0,047 | 0,041 | (0.0179, 0.1963) | 0,694 | 1,000 | Monte Carlo sample label permutation | 10000 |
| PostHoc Inter (Regional) | Mucoid vs PAO1 (Bottom) | nematic_order | 11 | 9 | - | W2 | 0,125 | 0,125 | (0.0182, 0.2938) | 0,201 | 0,804 | Monte Carlo sample label permutation | 10000 |
| PostHoc Inter (Regional) | Mucoid vs PAO1 (Bulk) | nematic_order | 11 | 9 | - | W2 | 0,056 | 0,052 | (0.0173, 0.2204) | 0,583 | 1,000 | Monte Carlo sample label permutation | 10000 |
| PostHoc Inter (Regional) | Mucoid vs PAO1 (Top) | nematic_order | 11 | 9 | - | W2 | 0,099 | 0,098 | (0.0164, 0.2636) | 0,337 | 1,000 | Monte Carlo sample label permutation | 10000 |

*Orientation Correlation ($G_2(r)$)*

$G_2(r)$ measures how far the alignment persists across the biofilm. It calculates the correlation between the orientations of pairs of bacteria separated by a distance $r$ (**equation 16**). It uses the second Legendre polynomial $P_2(x)$ to respect the head-tail symmetry:

$$G_2(r) = \langle \frac{3(\boldsymbol{u}_i \cdot \boldsymbol{u}_j)^2 - 1}{2} \rangle_{|\boldsymbol{r}_{ij}| \approx r} \qquad (16)$$

A slow decay of $G_2(r)$ indicates large domains of aligned bacteria (nematic order), while a fast decay to 0 indicates that alignment is only local. We calculated $G_2(r)$ for a 15 µm radius around each cell. All analyses were performed using a custom Python script.

**Supplementary Figures**

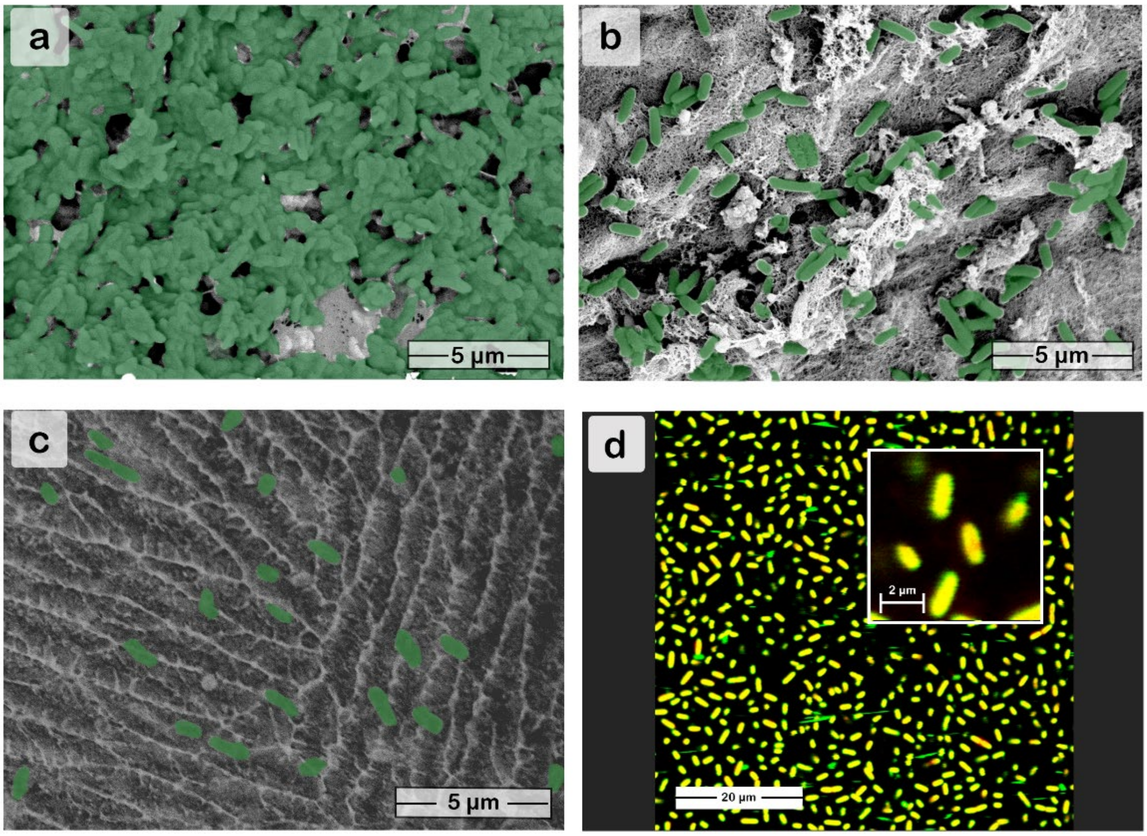


**Figure A4**: Effect of sample preparation on the spatial distribution of cells in 4-day-old mucoid PA biofilms. a) air-dried biofilm, b) critical-point dried biofilm, c) high-pressure frozen biofilm, and d) Syto9/PI-stained biofilm imaged by CLSM. The bacterial cells in a – c have been false-colored green for visibility.

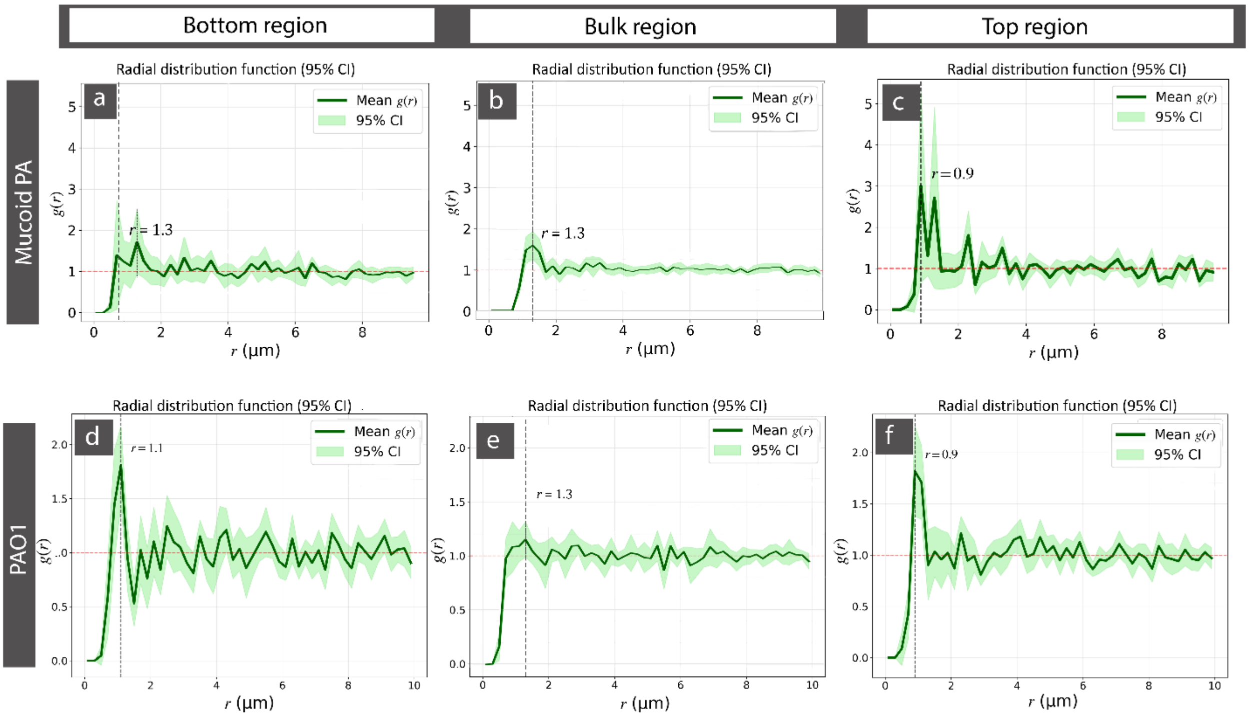


**Figure A5:** Averaged radial distribution function ($RDF$) analysis of PA biofilm spatial organization in cryo-SEM biofilm images, illustrating bacterial density as a function of radial distance from each cell center. a – c) $RDF$ from the bottom, bulk, and top regions of mucoid PA biofilms, respectively. d – f) $RDF$ from the bottom, bulk, and top regions of PAO1 biofilms, respectively. A total of ≥10 images per region were selected for calculating the average $RDF$. The green line represents the mean $RDF$, while the shaded area indicates the 95% confidence interval (CI). The black dotted line represents the most probable distance at which a cell finds a neighbor.

*RDF analysis*

In contrast to simple density measurements or the $NND$, the $RDF$ captures whether cells are randomly distributed or arranged with characteristic spacings. A value of $g(r) = 1$ corresponds to a random (Poisson) distribution of bacteria with the same overall density as observed in the sample, whereas pronounced peaks indicate preferred intercellular spacings. $RDF$ analysis is widely used in soft matter physics to study colloidal systems.[6,7]

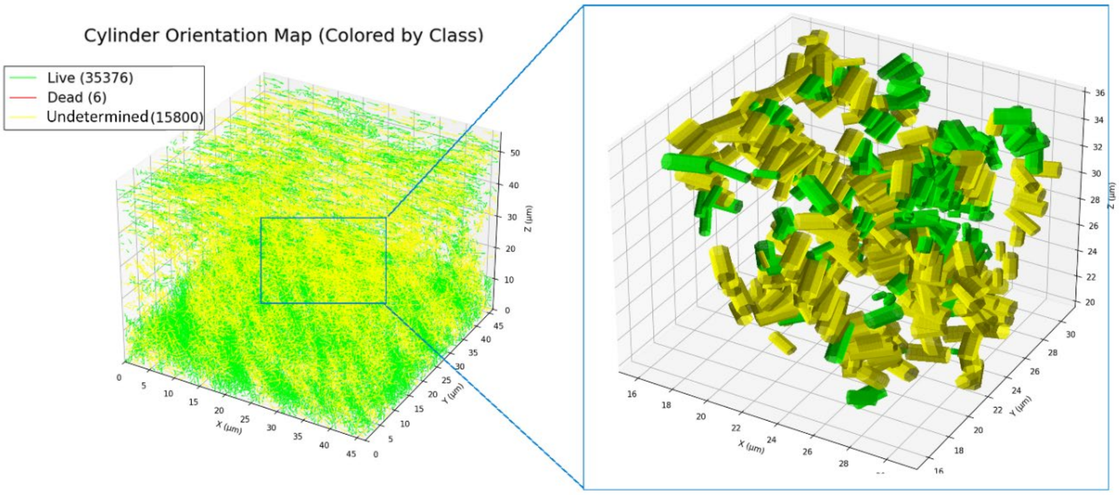


**Figure A6:** Three-dimensional orientation map of bacterial cells within the PA biofilm. Individual bacteria are represented as cylinders positioned and colored according to viability class (live (predominantly green), dead (predominantly red), and undetermined (strongly green and red)). The visualization highlights the spatial organization and orientation of cells across the biofilm volume.

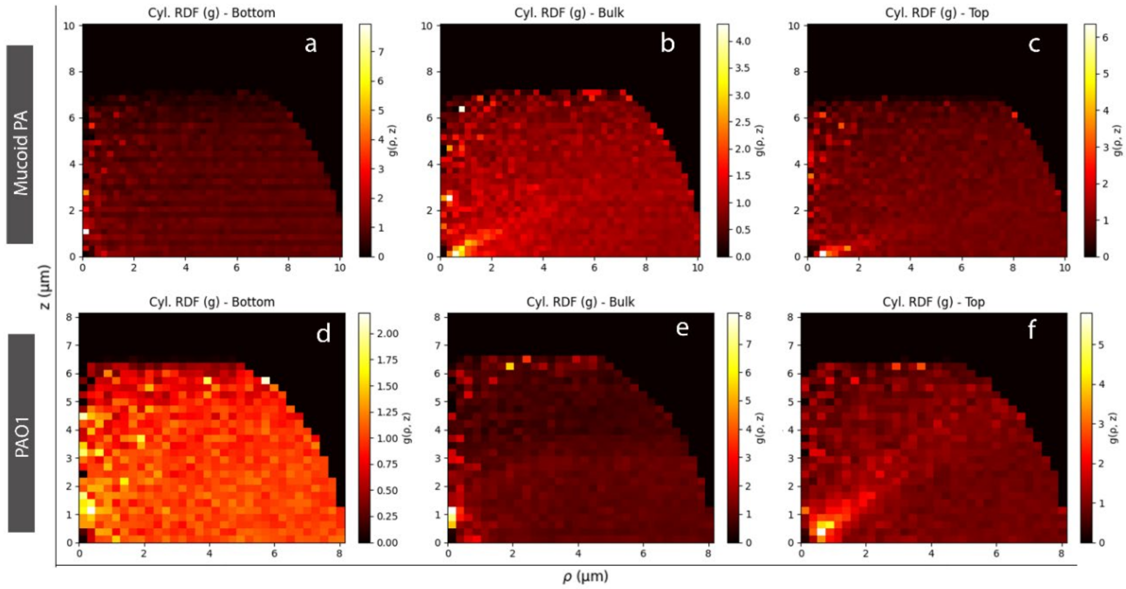


**Figure A7**: Representative 3D RDF analysis of PA biofilms across biofilm regions from CLSM data. Heatmaps were used to represent the RDFs of a-c) mucoid PA and d-f) PAO1 across the bottom, bulk, and top regions of the biofilms.

To match the $RDF$ of CLSM images to that obtained from the cryo-SEM images (**Figure A5**), only $g(r, z = 0)$ was compared (**Figure A7**). The $g(r, z = 0)$ analysis provided qualitatively the same results as the cryo-SEM $g(r)$, but with the most likely inter-bacteria distance several hundred nm shorter than in the cryo-SEM RDF analysis. The systematically shorter preferred distances suggest slightly closer cell distances but not dense clustering. This is likely attributable to methodological differences between the two imaging approaches. CLSM data are limited by diffraction-limited resolution, necessitating algorithmic distinction between closely spaced bacteria. This can misidentify a single large, diffuse bacterium as two, or more likely, position two closely spaced bacteria closer together than their actual distance because of their overlapping point-spread functions. Such errors will underestimate true average intercellular distances. In contrast, cryo-SEM provides superior resolution, easily enabling us to distinguish individual bacteria.

In the mucoid PA strains, the average first-peak values for $g(r, z = 0)$ were approximately 0.85 ± 0.13 μm (bottom region), 0.742 ± 0.07 μm (bulk region), and 0.746 ± 0.09 μm (top region), respectively, indicating a relatively consistent preferred spacing across the biofilm depth. In PAO1 biofilms, the corresponding average peak positions were approximately 1.11 ± 0.2 μm (bottom), 0.86 ± 0.09 μm (bulk), and 0.89 ± 0.14 μm (top), respectively. A significant ($p \leq 0.05$) difference in peak position between the bottom and the bulk was only found for PAO1. The larger confidence intervals in the bottom and top regions of the biofilms for both strains indicate greater heterogeneity in spacing for these regions across the imaged samples. The systematically shorter preferred distances obtained from CLSM than cryo-SEM suggest slightly closer cell packing in the fluorescence-based measurements. This is likely attributable to methodological differences between the two imaging approaches. CLSM data are limited by diffraction-limited resolution, necessitating algorithmic distinction between closely spaced bacteria. This can misidentify a single large, diffuse bacterium as two, or more likely, position two closely spaced bacteria closer together than their actual distance because of their overlapping point-spread functions. Such errors will underestimate true average intercellular distances. In contrast, cryo-SEM provides superior resolution, easily enabling us to distinguish individual bacteria.

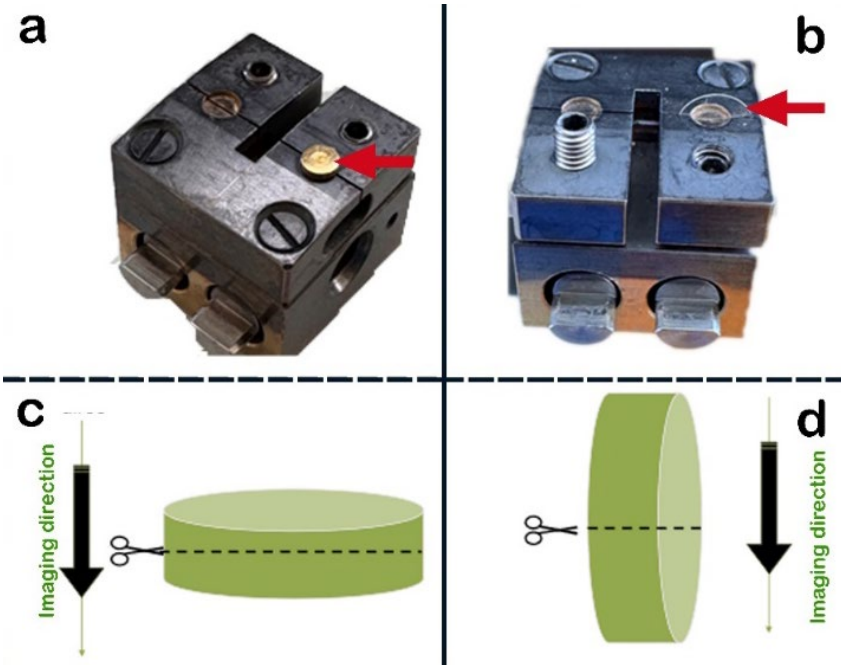


**Figure A8:** Images of sample holders with a) mounted carrier and b) a sapphire disc, highlighted with red arrows, and an illustration of sample fracture and imaging direction of biofilms grown on c) carriers and d) sapphire discs for cryo-SEM.

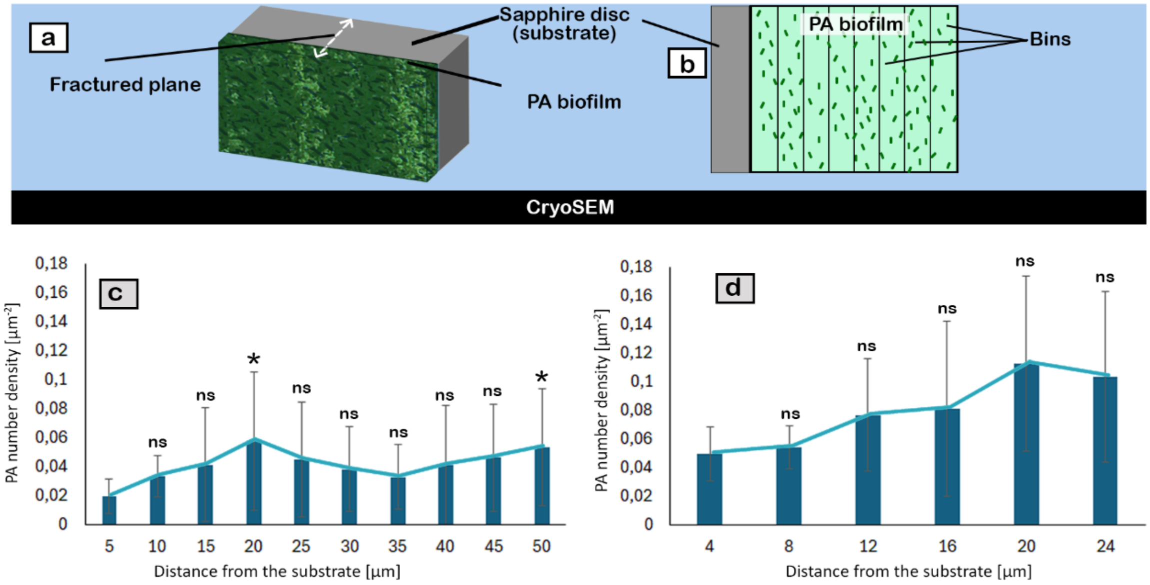


**Figure A9**: Spatial density profiles of PA biofilms. a) *P. aeruginosa* strains (mucoid and PAO1) were grown on sapphire discs for 4 days and subsequently high-pressure cryo-frozen and b) fractured orthogonally to the substrate to reveal a biofilm cross-section for spatial quantification. Cell density profiles for c) mucoid PA biofilms and d) PAO1 biofilms were generated. The first bars in the cell-density profiles of mucoid PA (5 µm) and PAO1 (4 µm), respectively, represent the density of cells closest to the substrate (sapphire disc) and were used as controls for statistical comparisons. Statistical significance was assessed relative to the control. * = $p \leq 0.05$; ns = $p \geq 0.05$

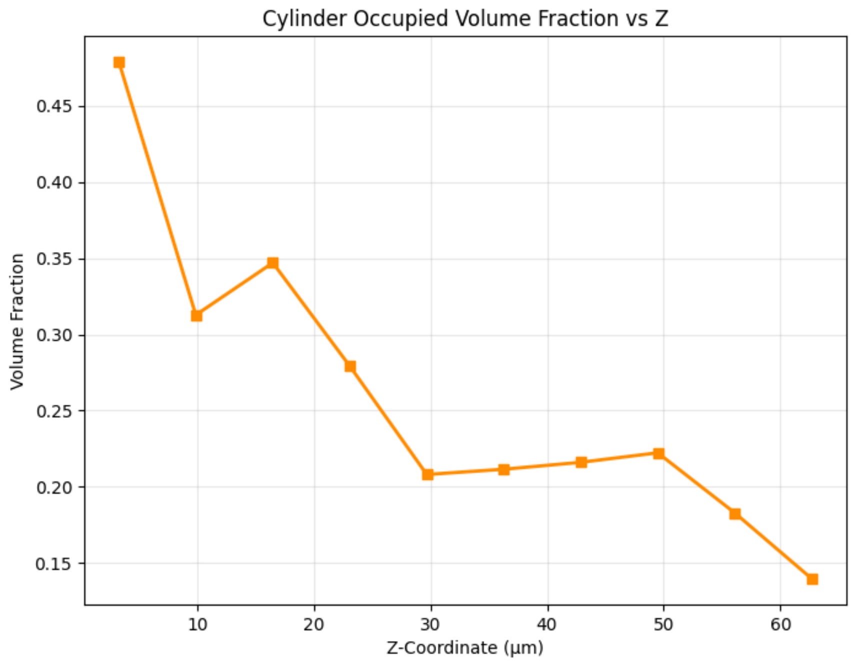


**Figure A10**: Cylinder (bacterial) volume fraction across biofilm depth for 4-day old mucoid PA biofilm

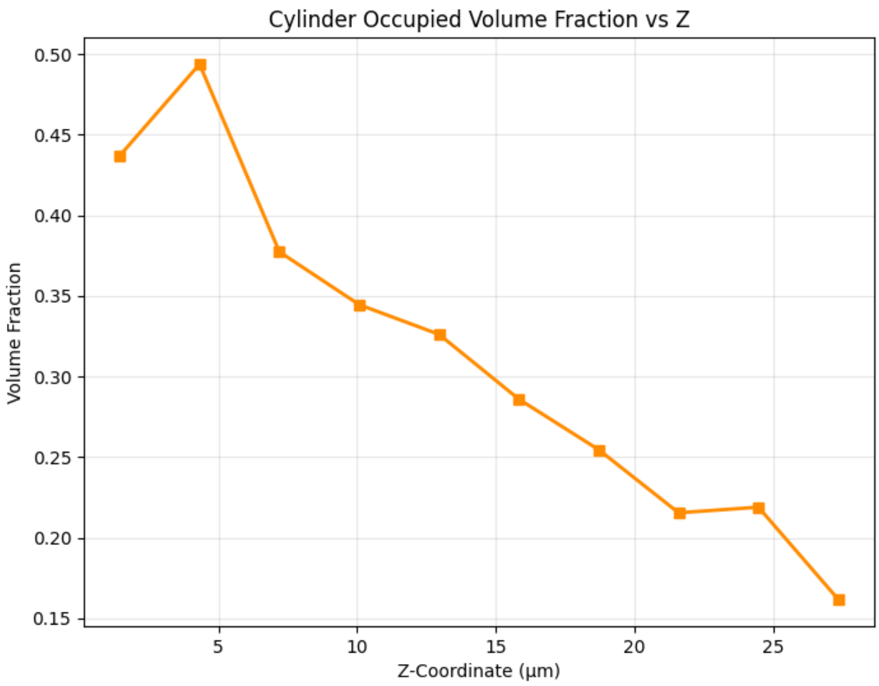


**Figure A11**: Cylinder (bacterial) volume fraction across biofilm depth for 4-day old PAO1 biofilm

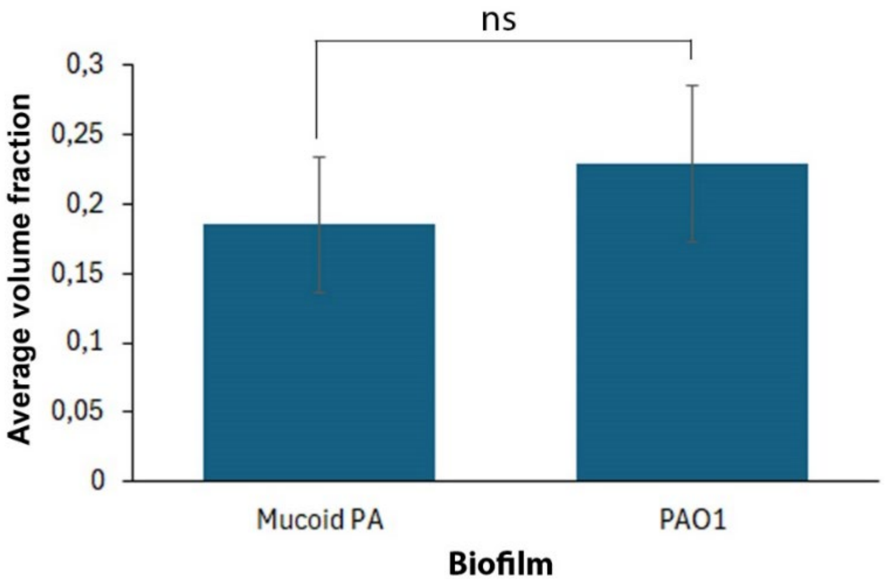


**Figure A12**: Comparison of biomass in PA biofilms. The chart shows the total biomass per area in mucoid and PAO1 strains. More than 6 independent samples were analyzed, and data presented as mean ± 95% CI. The statistical comparison between groups is indicated above the plot ($ns$ = not significant; $p \geq 0.05$).

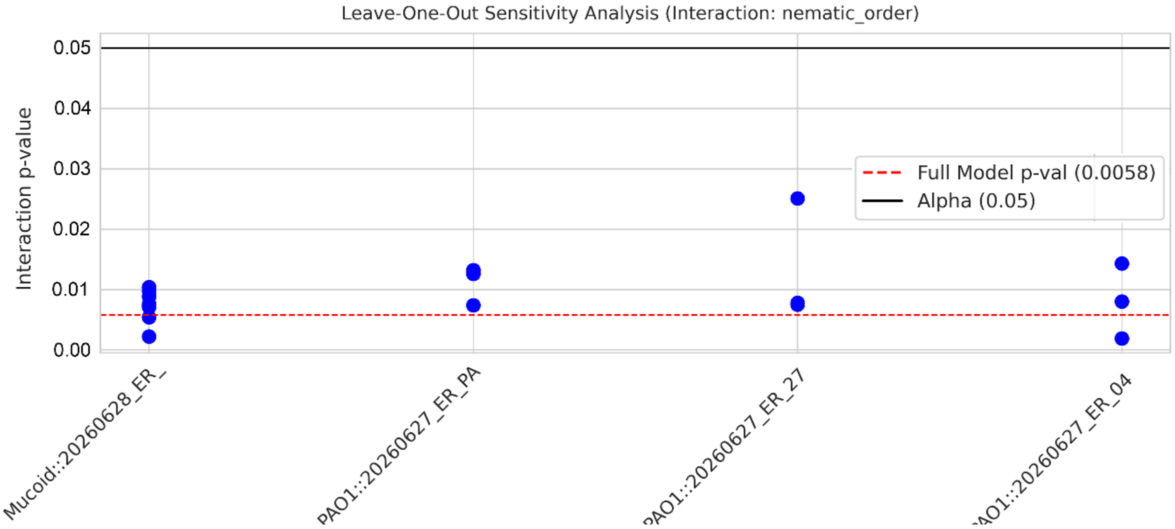


**Figure A13**: Leave-one-biofilm-out sensitivity analysis of the nematic-order interaction. The biofilm-type-by-region interaction test was repeated after excluding each biological biofilm in turn. The interaction remained below the significance threshold after removal of any single biofilm, indicating that the result was not driven by one individual sample.

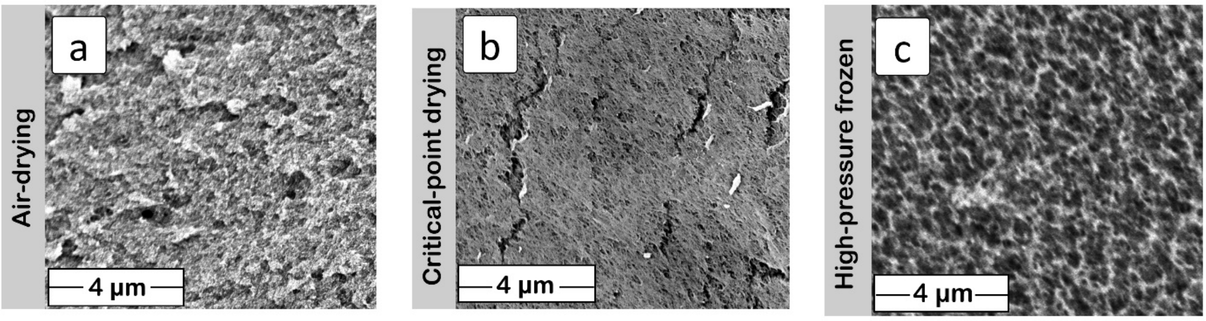


**Figure A14**: SEM micrographs depicting cross-sections of 0.5 wt% alginate hydrogels. Samples were prepared for imaging using a) air-drying, b) critical-point drying, and c) high-pressure freezing.

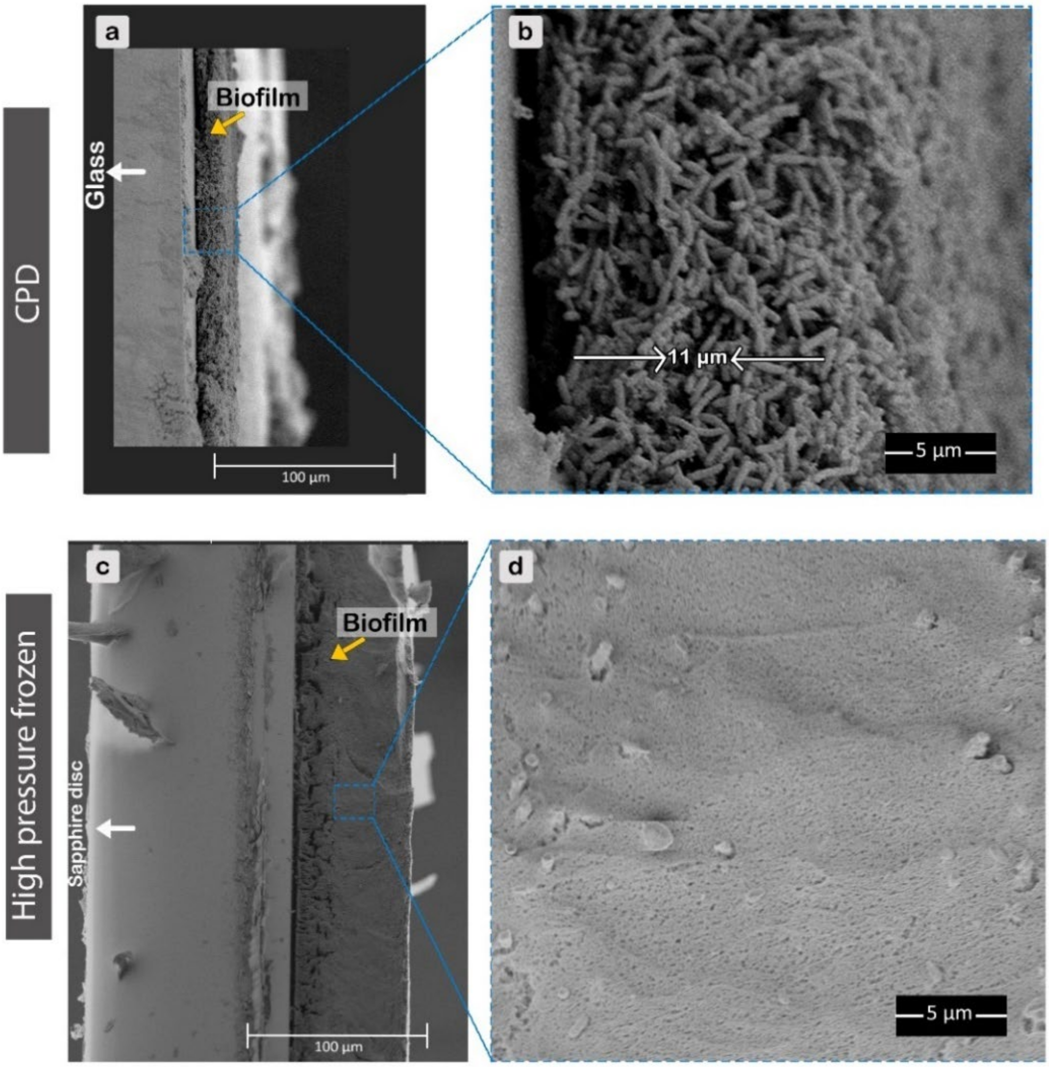


**Figure A15**: Micrographs of cross-sectioned mucoid PA biofilms prepared by different methods. Mucoid PA biofilms a) prepared by critical point-drying and b) prepared by cryo-freezing after 4 days of growth on cover glass and sapphire disc, respectively. Samples were afterwards fractured orthogonally to the substrate to reveal a biofilm cross-section.

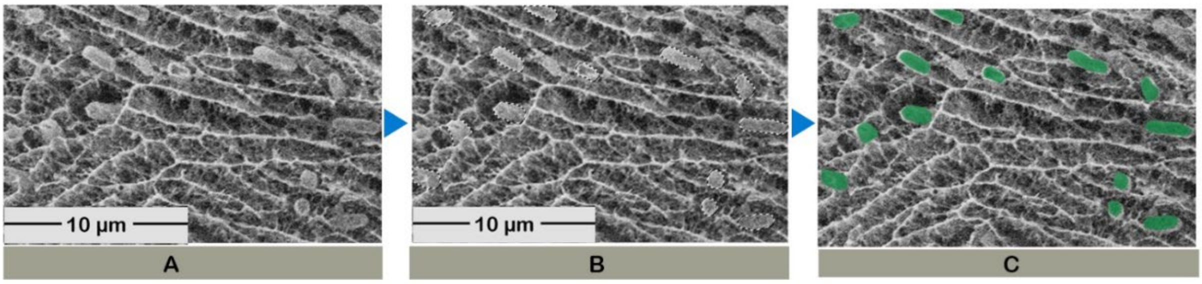


**Figure A16:** Workflow for cell coloring on SEM micrographs: A) representative SEM images of PA biofilms, B) manually masked bacterial cells, and C) masked bacterial cells after coloring

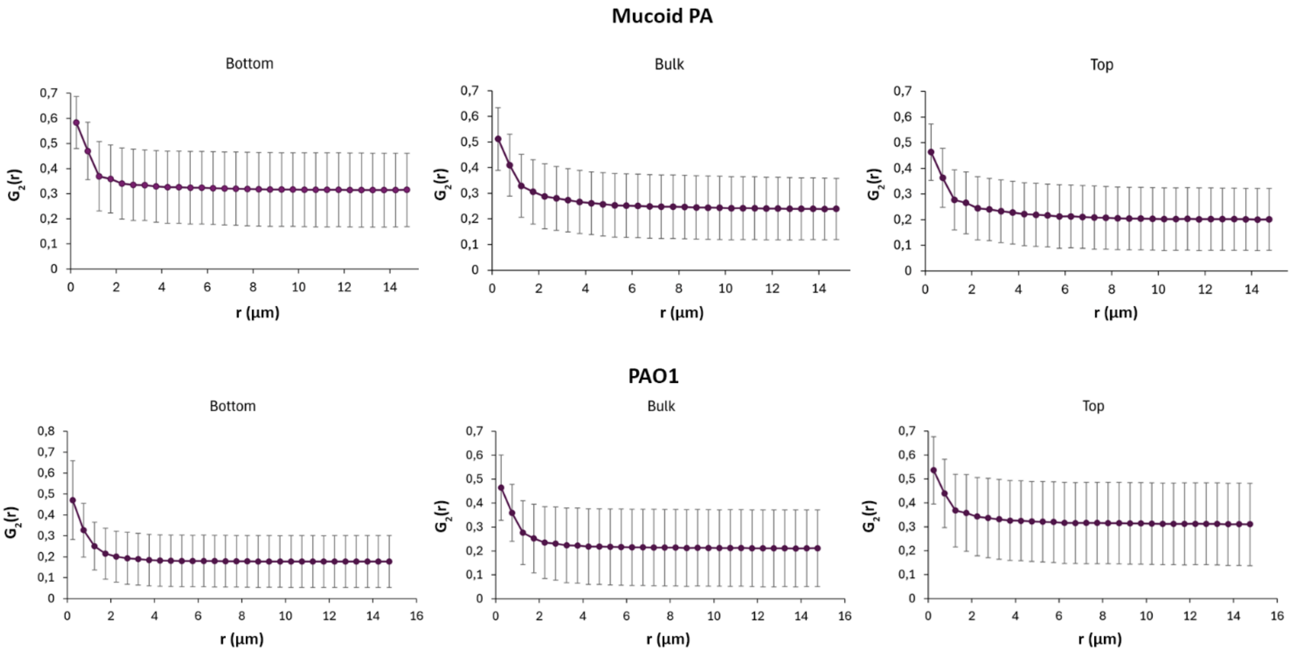


**Figure A17:** Averaged orientational correlation function with 95% confidence intervals calculated for the biofilm layers in 4-day-old mucoid PA and PAO1 biofilms

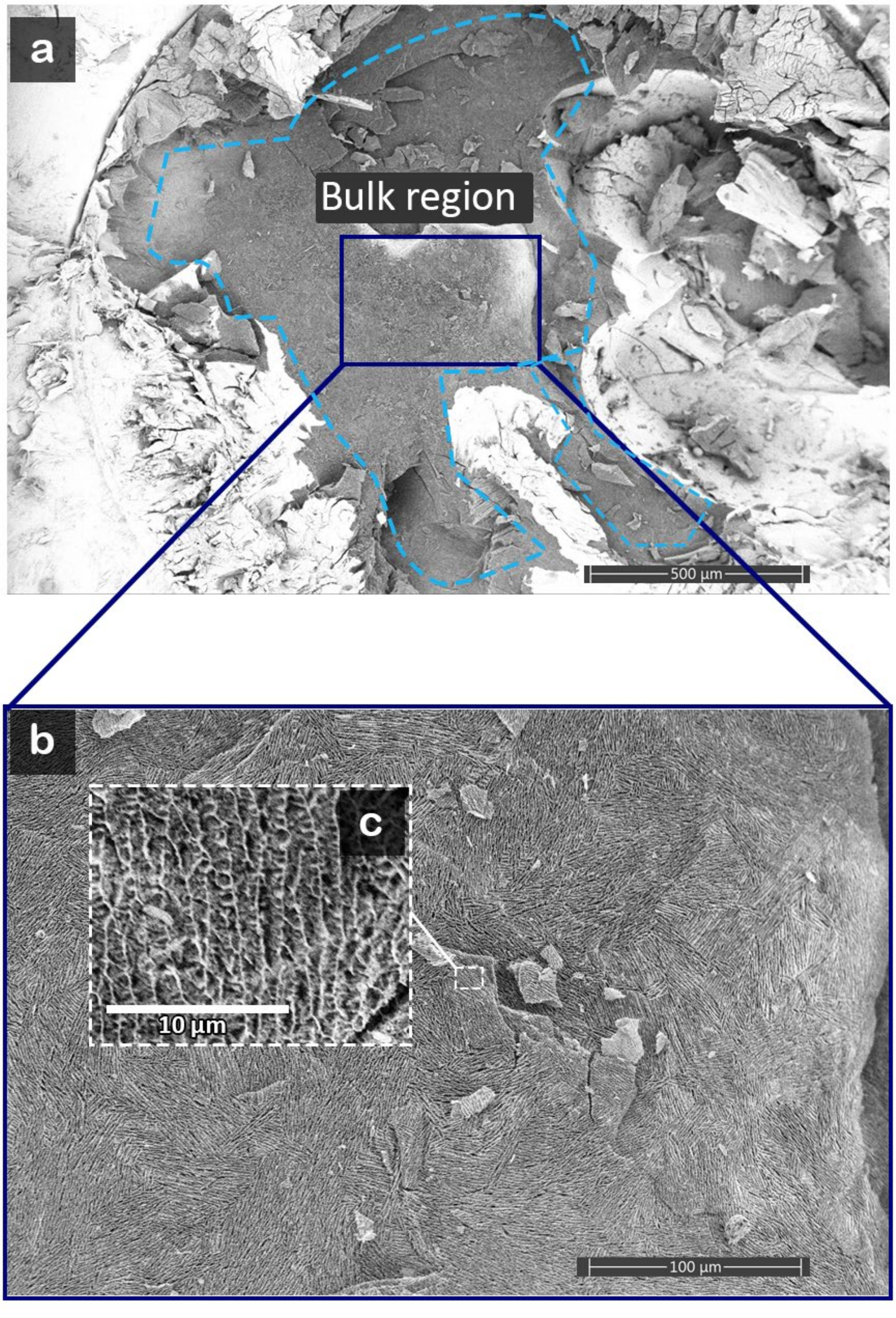


**Figure A18**: Cryo-SEM visualization of the bulk region of a mucoid PA biofilm. Cryo-fractured biofilm showing: a) low-magnification cryo-SEM overview showing the internal bulk region of the biofilm (outlined by the dashed blue line), highlighting the internal architecture away from the top and bottom regions. b) higher-magnification image of the boxed region in a), revealing ECM with pronounced fibrillar organization, and c) magnified inset illustrating the fine-scale fibrillar ECM ultrastructure within the bulk region.

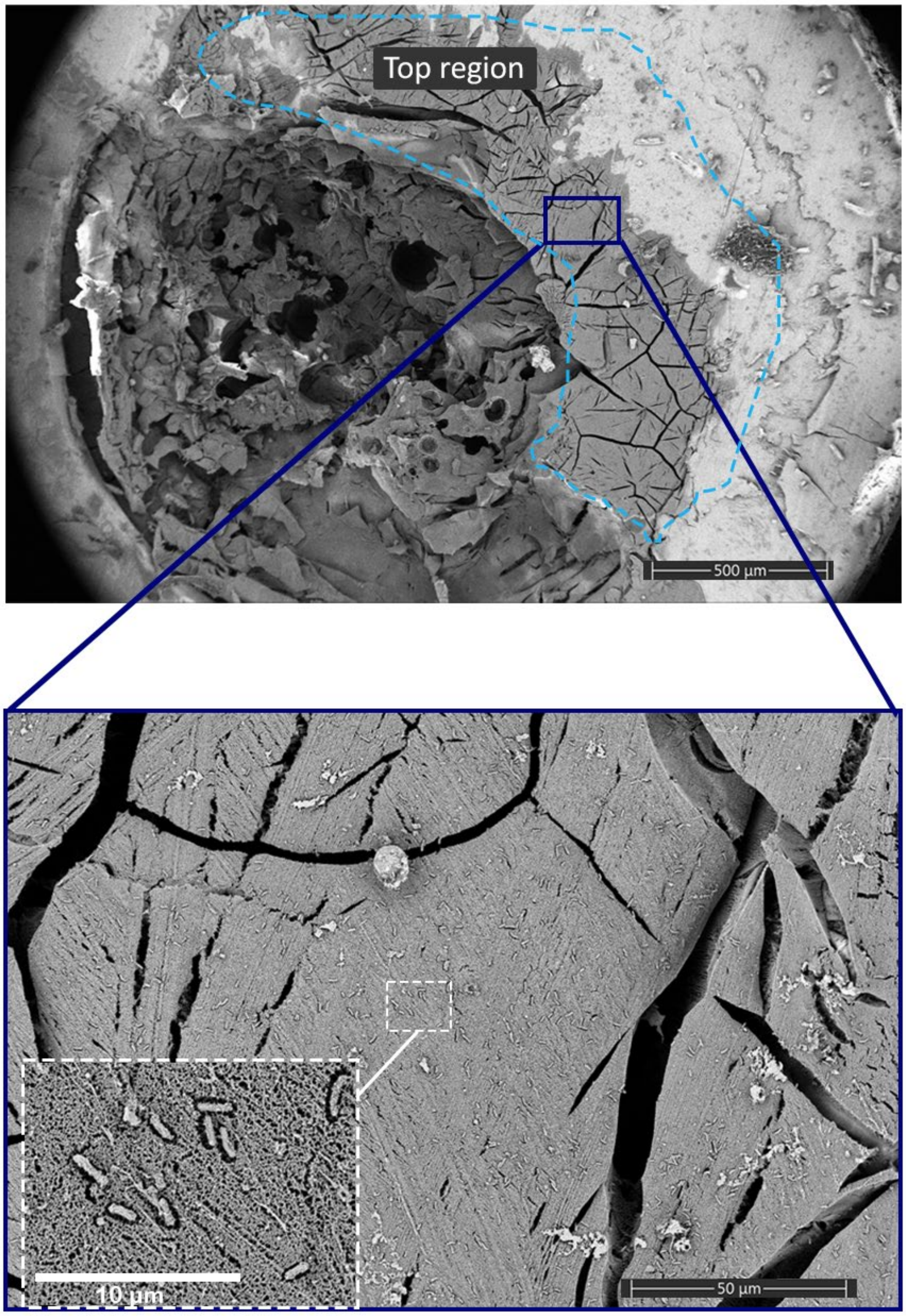


**Figure A19**: Cryo-SEM visualization of the top region of a mucoid PA biofilm. Cryo-fractured biofilm showing: a) low-magnification cryo-SEM overview showing the top region of the biofilm (outlined by the dashed blue line), b) higher-magnification image of the boxed region in (a), revealing ECM as a mesh with spatially organized cells, and c) magnified inset illustrating the fine-scale ECM ultrastructure at the top region.

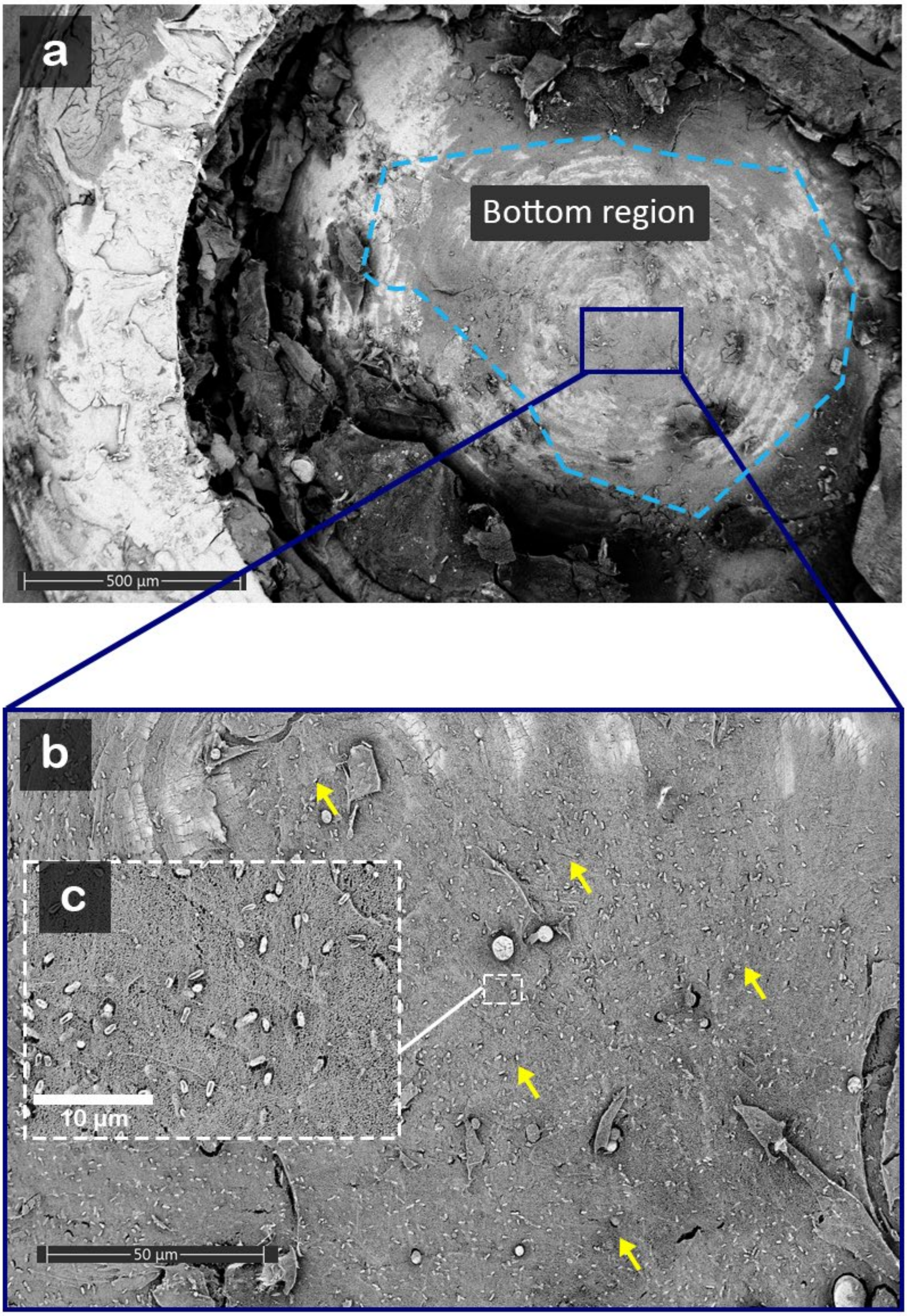


**Figure A20**: Cryo-SEM visualization of the bottom region of a mucoid PA biofilm. a) Low-magnification cryo-SEM image showing the fractured biofilm cross-section, with the edge and bottom of the sample carrier indicated. The white arrowheads mark the bulk region of the biofilm. b) Higher-magnification view of the boxed region, revealing bacterial cells embedded within a dense ECM in the substrate-proximal layer. Yellow arrows indicate bacterial cells within the matrix. c) Inset showing biofilm details at higher resolution.

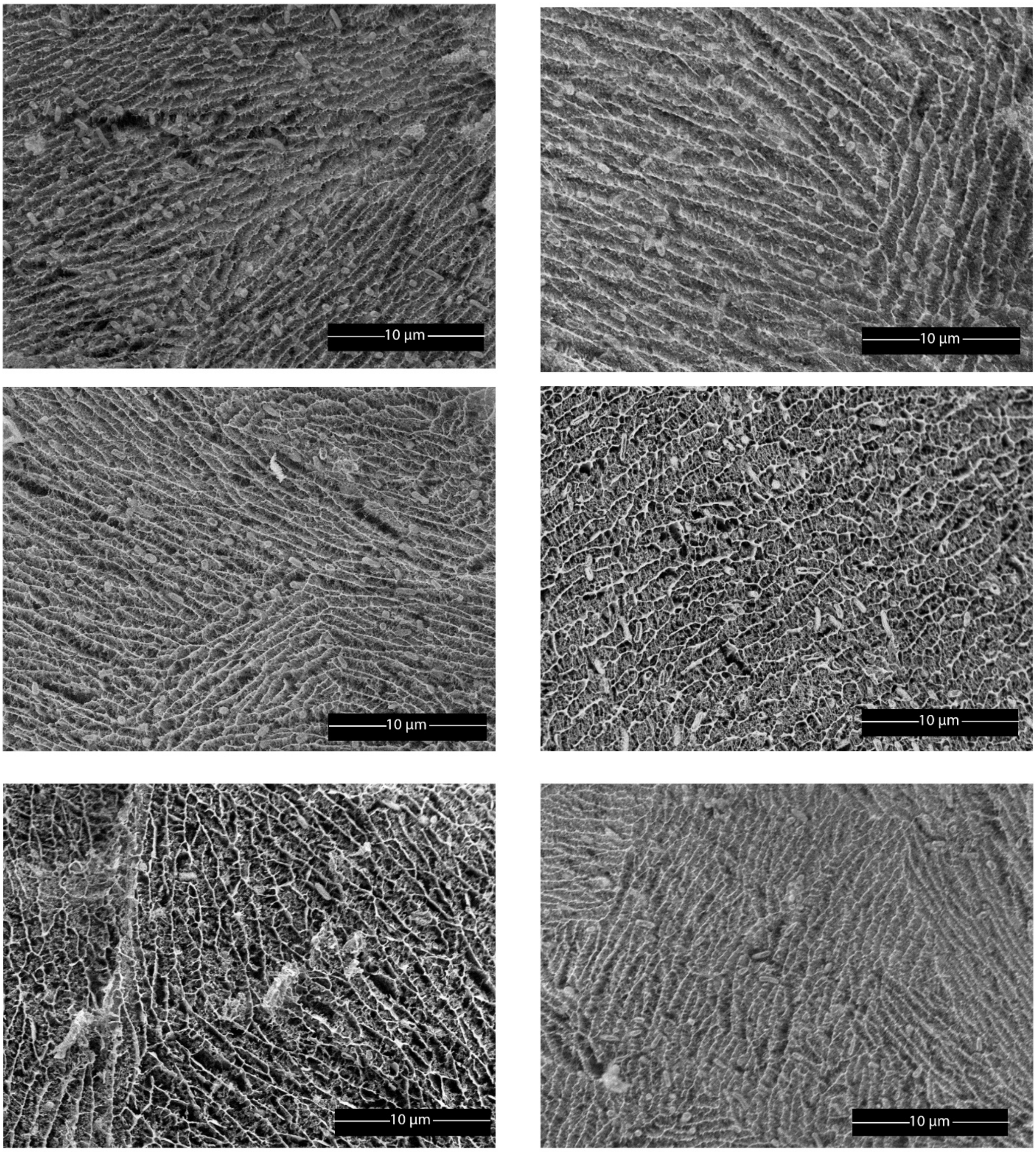


**Figure A21**: Cryo-SEM visualization of the bulk region of a mucoid PA biofilm from different samples, highlighting similar fibrillar organization.

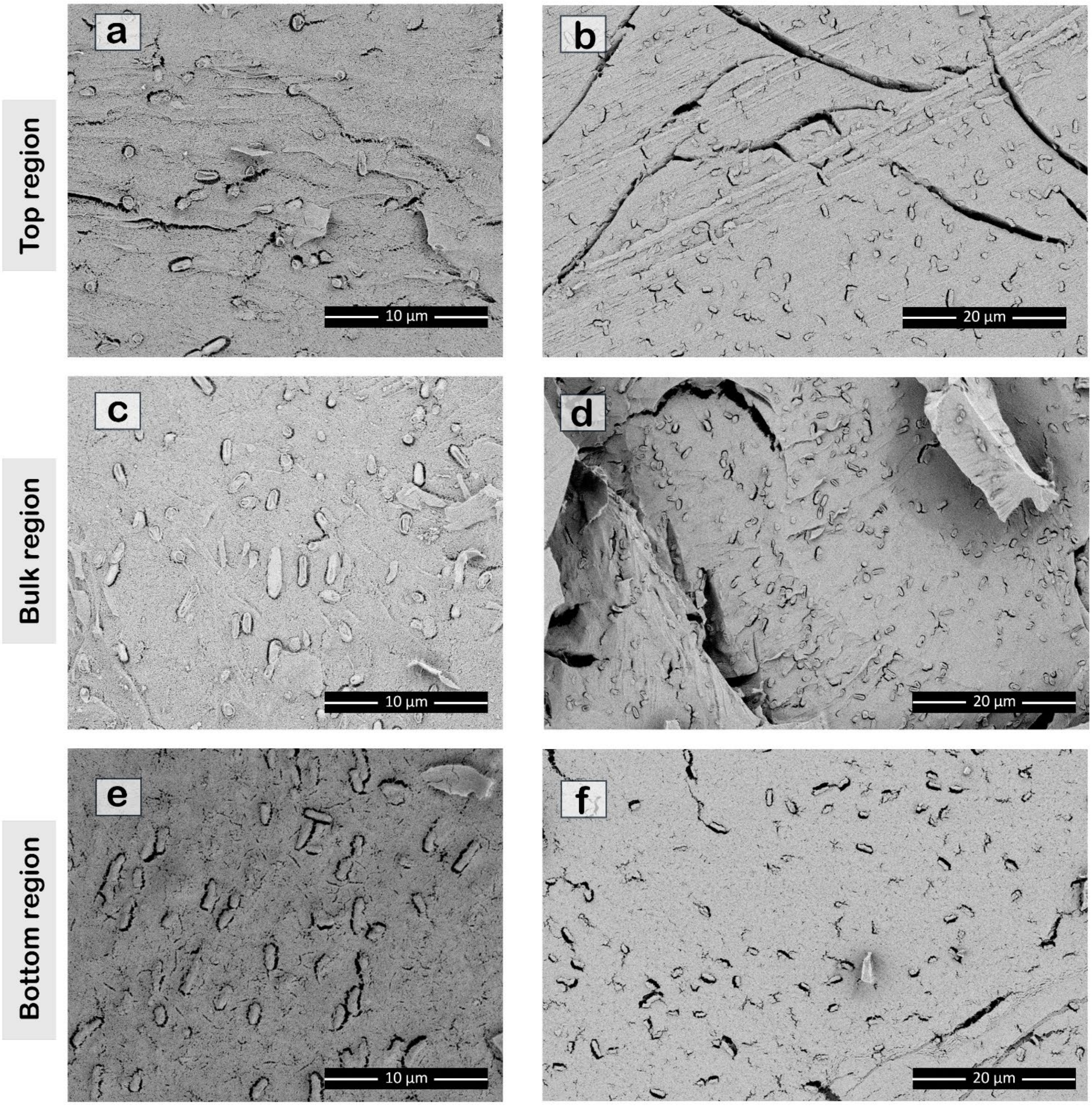


**Figure A22**: Gallery of images from high-pressure frozen PAO1 biofilms across different regions. a-b) top, c-d) bulk, and e-f) bottom regions of PAO1 biofilm from different samples.

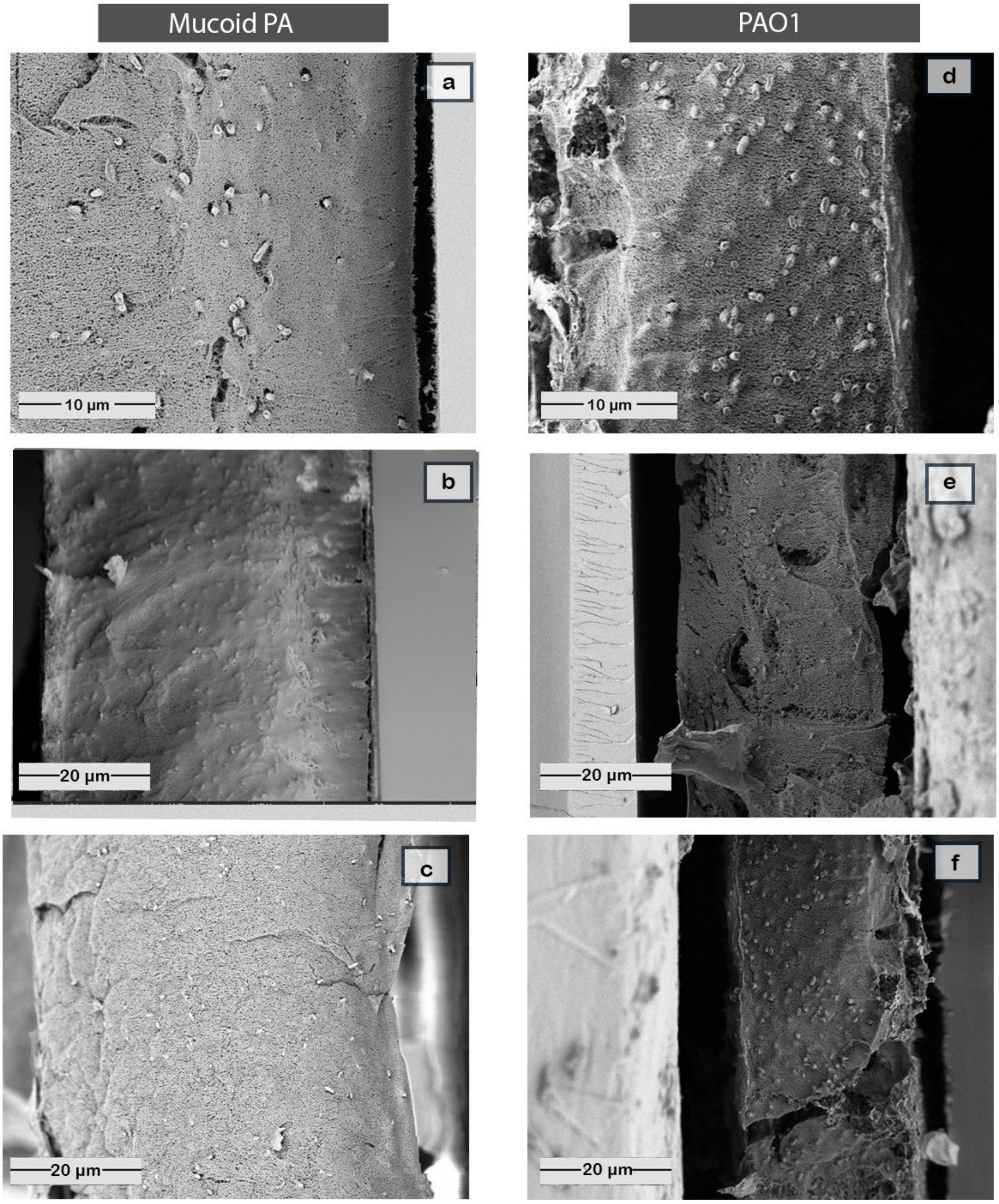


**Figure A23**: Gallery of cryo-SEM images from fractured a-c) mucoid PA and d-f) PAO1 biofilms grown on sapphire discs for 4 days.

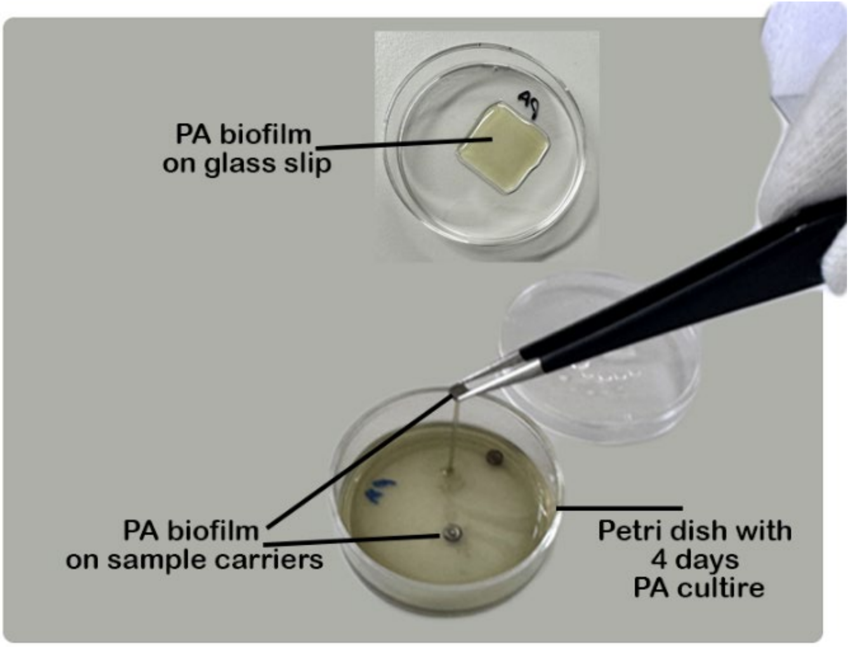


**Figure A24**: Representative images of PA biofilms grown for 4 days on different substrates. Biofilms formed on cover glass (top) and on sample carrier substrates (bottom) are shown in Petri dishes containing PA cultures. The image illustrates the experimental setup used for biofilm growth prior to sample processing and imaging.

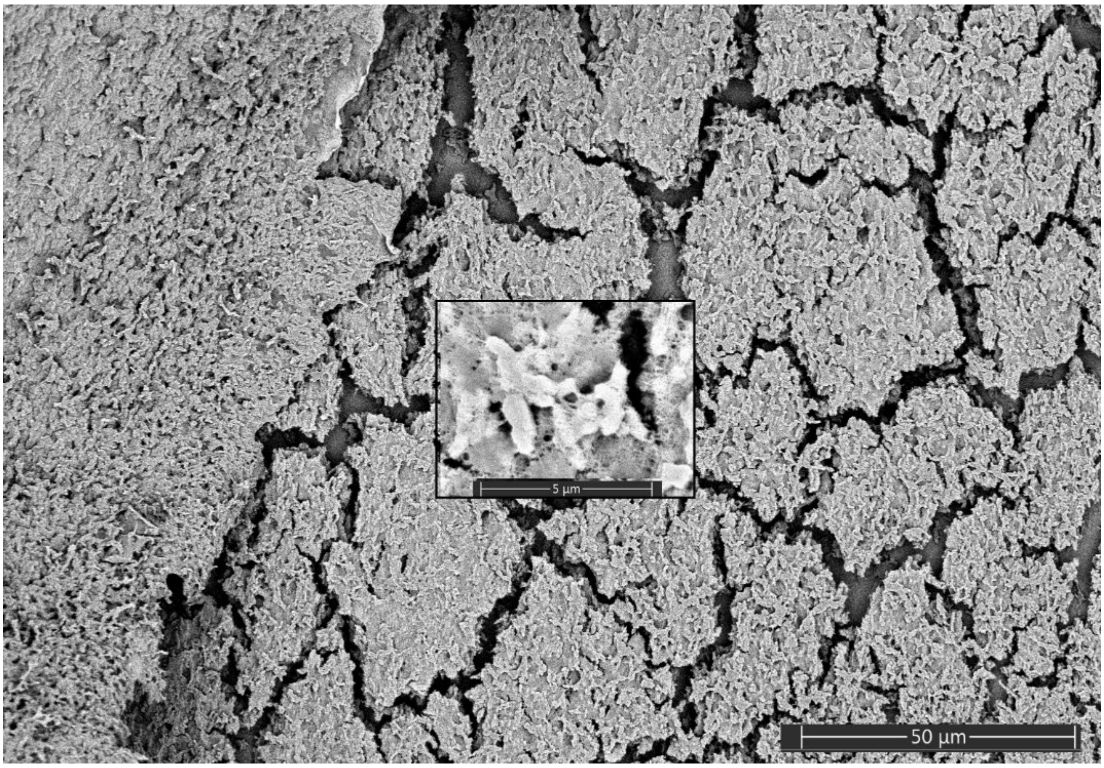


**Figure A25**: SEM micrographs of a 4-day-old mucoid PA biofilm prepared by air-drying.

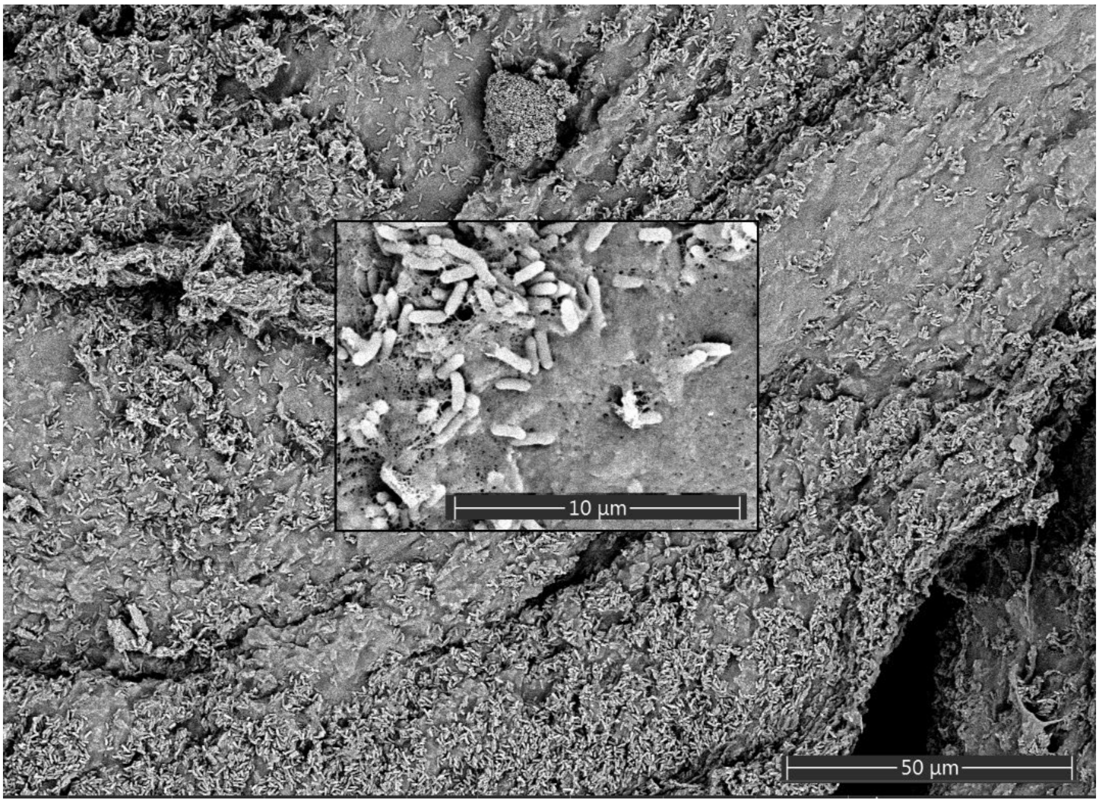


**Figure A26**: SEM micrographs of a 4-day-old mucoid PA biofilm prepared by critical-point drying.

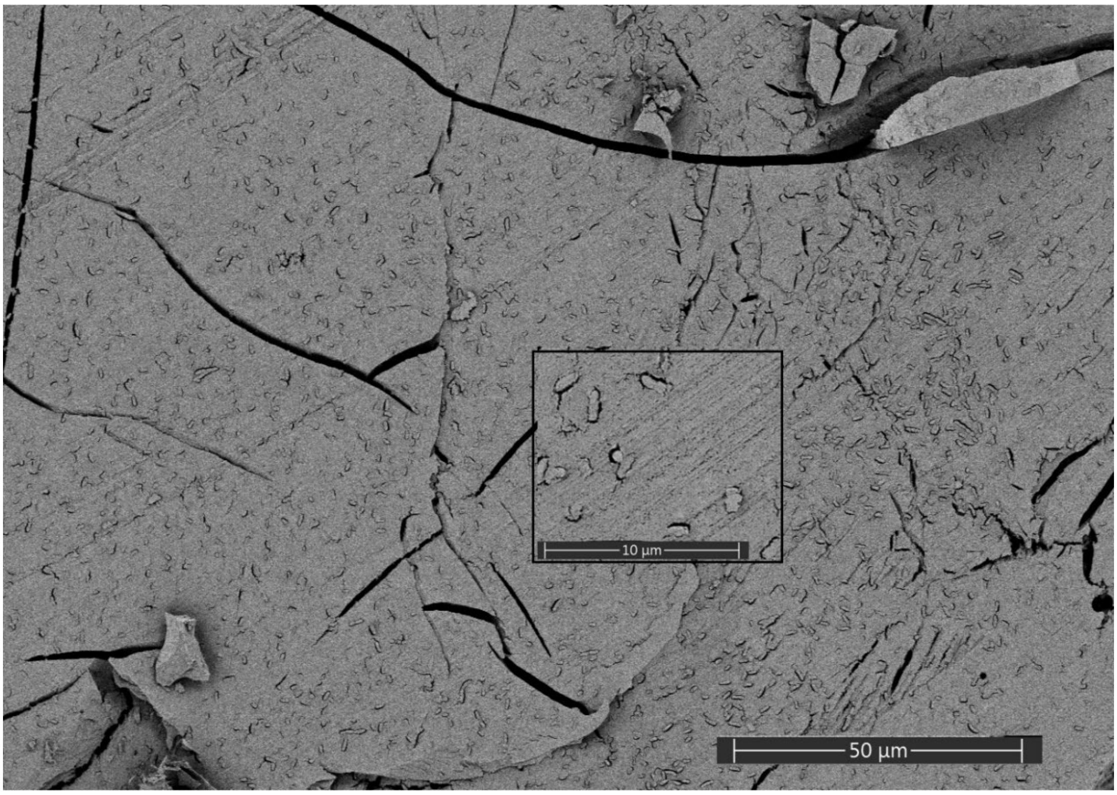


**Figure A27**: Cryo-SEM micrographs of the top region of a PAO1 biofilm.

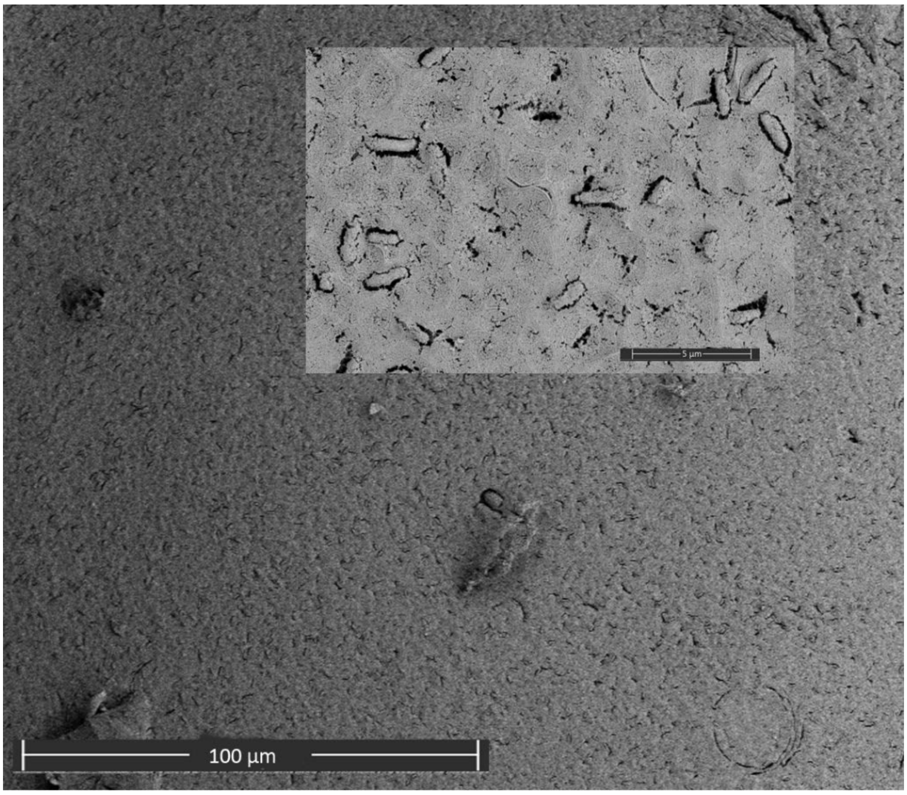


**Figure A28**: Cryo-SEM micrographs of the bottom region of a PAO1 biofilm.

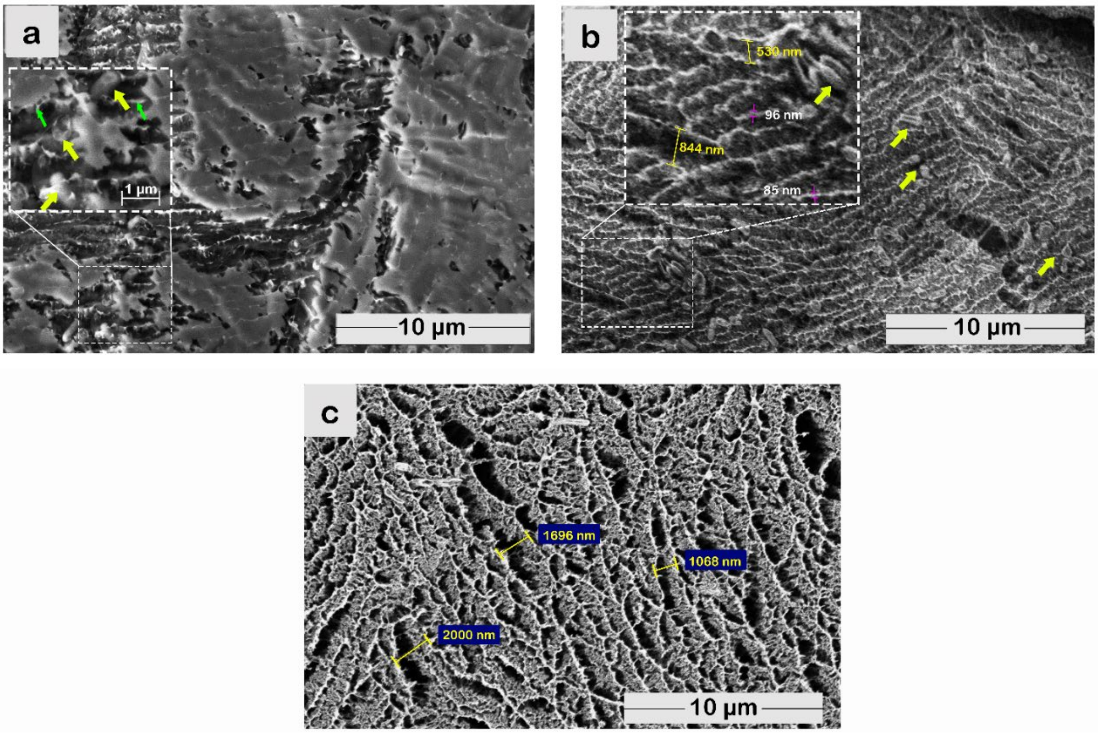


**Figure A29**: Cryo-SEM micrographs from the bulk region of 4-day-old mucoid PA biofilms. a) 0 s etching, b) 20 s etching, and c) deep etching (60 s). The yellow arrows indicate the outlines of bacterial cells, while the green arrows indicate the contours of the ECM.
